# Dopamine-gated memory selection during slow wave sleep

**DOI:** 10.1101/2020.05.23.112375

**Authors:** Hanna Isotalus, Will J Carr, George G Averill, Oliver Radtke, James Selwood, Rachel Williams, Elizabeth Ford, Liz McCullagh, James McErlane, Cian O’Donnell, Claire Durant, Ullrich Bartsch, Matt W Jones, Carlos Muñoz-Neira, Alfie R Wearn, John P Grogan, Elizabeth J Coulthard

**Affiliations:** Clinical Neurosciences, Translational Health Sciences, Bristol Medical School, University of Bristol, Bristol; Digital Health, Faculty of Engineering, University of Bristol, Bristol; Southmead Hospital, North Bristol NHS Trust, Bristol; Experimental Psychology, University of Bristol, Bristol; School of Physiology, Pharmacology and Neuroscience, University of Bristol; Department of Neurology, Centre for Movement Disorders and Neuromodulation, Medical Faculty, Heinrich-Heine-University, Düsseldorf.; Production Pharmacy, Bristol Royal Infirmary, University Hospitals Bristol and Weston NHS Trust; School of Computer Science, Electrical and Electronic Engineering, and Engineering Mathematics, University of Bristol, Bristol; Nuffield Department of Clinical Neurosciences, University of Oxford, Oxford

## Abstract

The human brain selectively stores knowledge of the world to optimise future behaviour, automatically rehearsing, contextualising or discarding information to create a robust record of experiences. Storage or forgetting evolves over time, particularly during sleep. We have previously shown that dopamine given in the form of L-DOPA tablets improves long-term memory in Parkinson’s disease, but only when given overnight. L-DOPA is already prescribed widely with a good safety profile and could potentially be rapidly repurposed to improve cognitive performance and improve quality of life in, for example, early Alzheimer’s Disease, if we understood the best time of day to prescribe. Therefore, we sought to test how dopamine shaped long-term memory formation before and during sleep in a double-blind randomised placebo-controlled cross-over trial of healthy older adults (n = 35). We administered L-DOPA after word-list learning to be active during repeat exposure to a proportion of the words and during subsequent nocturnal sleep. Nocturnal dopamine accelerated forgetting for words presented once but it did not affect memory for words presented twice. During slow wave sleep, L-DOPA also increased spindle amplitude around slow oscillation peaks. Larger dopamine-induced difference in word memory was associated with a larger increase in spindle amplitude. Dopamine-dependent memory processing may therefore modulate spindles dependent on slow-oscillation phase. Further, overnight dopamine increased total slow wave sleep duration by approximately 11%. This pharmaceutical modification of slow wave sleep may have potential health-enhancing benefits in old age that could include cognitive enhancement and Alzheimer’s prevention.

**One Sentence Summary:** Dopamine before sleep promotes forgetting of weak memory traces associated with increased spindle amplitude around the peak of a slow oscillations.

## Introduction

The brain selectively extracts and stores important details of our daily lives, while demoting irrelevant information - you have probably forgotten where you parked your car while shopping last week, but you will remember your parking slot in an airport carpark after a week’s holiday. When memories are first encoded they form traces known as engrams – or changes in neuronal and synaptic activity that represent a memory (*1, 2*). Depending on context and relevance, engrams can be integrated within memory networks for the long term or forgotten through a set of processes that start immediately and progress during wake and sleep (*3–5*).

The complex milieu that underpins memory formation depends on several neurotransmitters including dopamine. This transmitter is released from two midbrain areas - locus coeruleus and ventral tegmental area – which directly project to the hippocampus (*6*). Exogenous dopamine administration can modulate memory persistence, particularly after initial learning (*7–10*). In humans with dopamine depletion due to Parkinson’s disease, memory consolidation improves with overnight administration of L-DOPA (Levodopa – which increases dopamine concentrations in the brain), but the timing of the dopamine manipulation relative to learning critically determines its effects on memory (*8, 11*).

During memory encoding and shortly after, engrams of important information can be prioritised for storage, based either on previous knowledge, repeated exposure, or other associations, such as financial or emotional value (*12, 13*). At a molecular level, synaptic, ‘tagging’ for information prioritised for storage, protein synthesis and synaptic modifications occur within hours of encountering information (*14, 15*). The dopaminergic connection between midbrain and hippocampus may selectively bias long-term memory storage by altering synaptic tagging or the protein synthesis involved in synaptic tagging (*15–17*).

As well as prospectively prioritising, or tagging, memories for later replay during sleep, dopamine may directly act during sleep *per se* (*18*). As engram storage evolves, newly acquired memories are spontaneously repeated (*17*); sleep affords an optimal neurophysiological state during which to enact this process - although replay occurs during wake too (*19*). Patterns of activation within hippocampal neuronal assemblies are selectively replayed during sharp wave ripples which are, in turn, temporally coupled to sleep spindles and slow oscillations, prominent during non-REM sleep (*20–24*).

Contextual information at a later time-point can also retroactively alter the likelihood that previous memories are stored for longer term (*25–27*). Dopaminergic modulation of memory may underpin contextually-driven modification of engrams (*16, 28–30*). Exogenous administration of dopamine may therefore alter the likelihood of memory stabilisation when given *after* initial learning, during re-exposure and consolidation.

It is important to point out that as well as actively prioritising relevant memories for storage, several neurobiological substrates could promote forgetting (*31, 32*). In drosophila, the ‘forgetting cells’ promote changes in cellular signalling that weaken the engram by releasing dopamine. Dopamine enhances encoding of new memories at the cost of triggering forgetting of competing information (*33, 34*). This dopamine-induced strategic forgetting is selective to weakly encoded memories – presumably, an automatic strategy for ensuring retention of more behaviourally relevant information. In humans, retroactive retrieval of already learnt information has been shown to simultaneously enhance memory for the retrieved information while inducing forgetting of contextually related information (*27*). This retrieval induced forgetting is caused by an *active* inhibition of the competing memory traces during recall (*35, 36*).

Here we propose a role for dopamine in *selecting* memories for long-term storage. Specifically, we propose that, after initial learning, dopamine biases human memory in favour of strongly encoded memory traces as opposed to weaker traces. We tested the hypothesis that L-DOPA given after initial learning (to be active during re-exposure and subsequent sleep) will increase memory retention compared to placebo. We also predicted that no such effect would be observed for words that were not re-exposed. We predicted the primary effects of dopamine during long-term memory evolution would be mediated through modulation of slow wave sleep duration and sleep spindle characteristics.

In this double-blind randomised within-subjects placebo-controlled trial, we show that a single dose of exogenous dopamine (L-DOPA) given after learning (to be active during re-exposure and subsequent nocturnal sleep) unexpectedly accelerates forgetting of non-repeated information effectively biasing memory selection away from weakly encoded items. Investigation of sleep characteristics further showed that spindle amplitude during slow wave sleep was higher on L-DOPA compared to placebo, and this effect was slow oscillation phase dependent. The magnitude of this increase in spindle amplitude correlated with the behavioural effect of dopamine on memory selection. Further, we show that nocturnal L-DOPA increases slow wave sleep duration by ~11%.

We also report a secondary placebo-controlled randomised control study in which we found that L-DOPA did not affect encoding or retrieval of learned words, further suggesting that the effects of L-DOPA in the main experiment were enacted during repetition, consolidation and/or sleep.

## Results

To study the relationships between dopamine, sleep and forgetting, we carefully timed administration of L-DOPA to increase dopamine concentration in healthy older adults across two placebo-controlled double-blind randomised crossover experiments: the main experiment and a secondary experiment. The overarching structure enabled targeting of L-DOPA to different memory processes – in the main experiment (**Fig. 1a.**), we explored the effects of dopamine on memory consolidation by administering long-acting L-DOPA after learning, to be active after initial learning and during nocturnal sleep (*37*). In the secondary experiment, we used short-acting L-DOPA to target memory retrieval (testing) or encoding (learning), but not sleep.

**Fig. 1.**
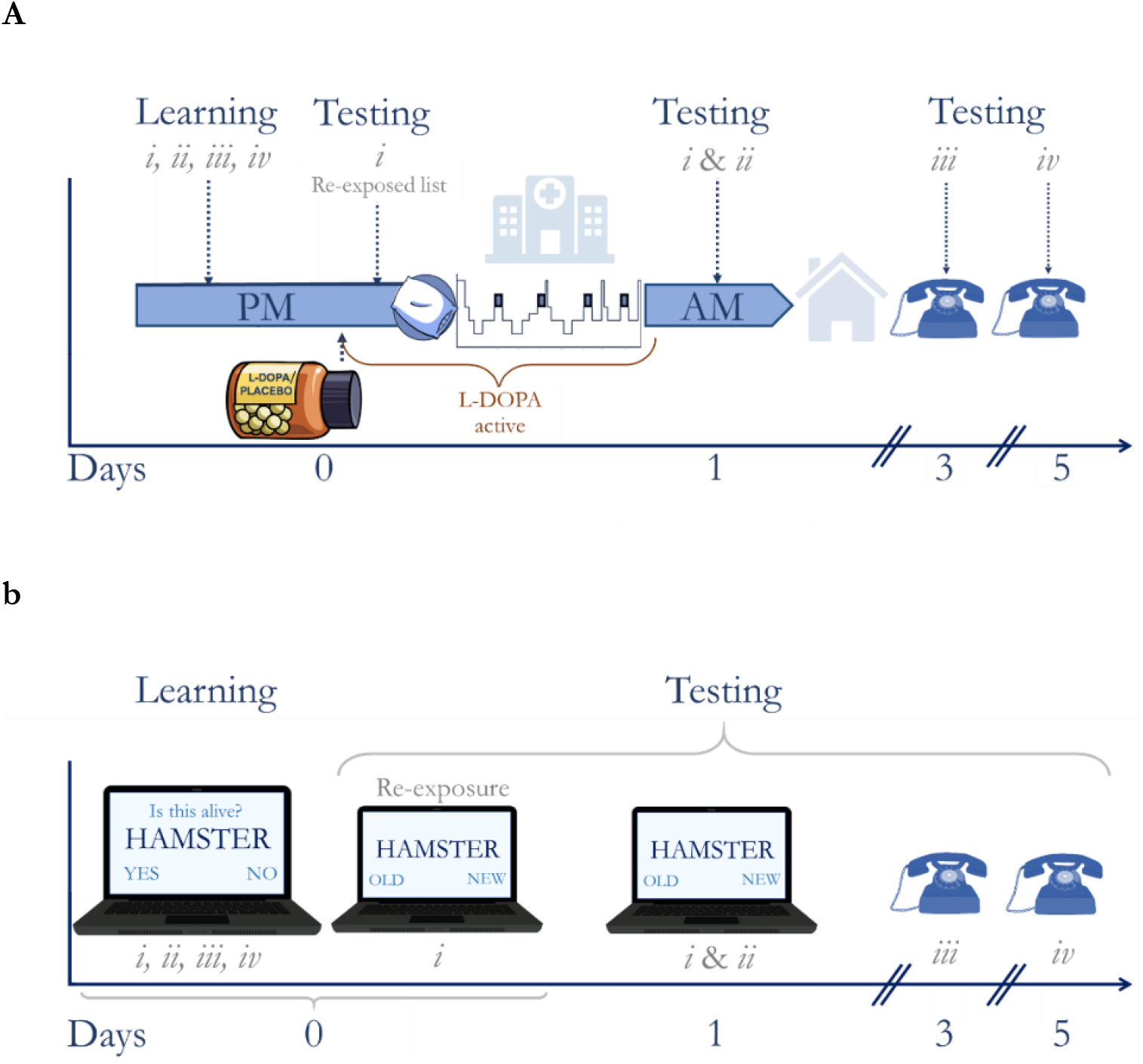
Experiment 1 Study procedure. **a**. In this placebo-controlled randomised crossover trial, healthy elderly volunteers completed two overnight sleep polysomnography visits. In the evenings, they learnt a set of words (**Fig. 1b.**) 45min *before* receiving 200mg L-DOPA CR or placebo. 75min *after* dosing a portion of the words were re-exposed. The orange bracket denotes when L-DOPA was active. Therefore, L-DOPA was active during re-exposure and sleep but not during learning or memory tests on days 1, 3, or 5. Apart from treatment (L-DOPA or placebo) the nights were identical. Each volunteer completed both nights. **b**. Participants were asked to memorise 80 words shown on a computer screen one at a time. The words were separated into four lists for testing (Lists *i, ii, iii, iv* – 20 words each) but during learning they were shown in a random, interleaved, order. 2h after learning, participants were re-exposed to List *i* by a recognition test. The following morning, ~12 hours after initial learning, memory for Lists *i* and *ii* was tested (random, interleaved), while lists *iii* and *iv* were tested 3 and 5 days later over the phone. Each test was performed using a recognition test with a unique set of distractor words. The testing procedure was fully explained to participants before learning.

In the main experiment, 35 healthy elderly volunteers (age = 68.9 ± 3.5 years; 22 Female) completed two overnight study visits (**Fig. 1a.**) which were identical except for treatment allocation. On each visit, we administered controlled release L-DOPA (CR; co-beneldopa 200/50mg) or placebo *after* participants had learnt information (word Lists *i, ii, iii and iv,* **Fig. 1b.**). Participants were re-exposed to a quarter of the items (List *i*) shortly after L-DOPA (or placebo) administration through a recognition memory test – this manipulation was performed to strengthen the memory for each List *i* word. Memory for the re-exposed items (List *i*) was tested the following day together with a matched number of items that had not been re-exposed (List *ii* – weak memory). Memory for the remainder of the items was probed 3 or 5 days after learning (Lists *iii* and i*v*). The participants knew some words would be tested both in the evening and in the morning, and the remainder of the words would only be tested once.

Initial learning occurred before L-DOPA (/ placebo), whereas memory re-exposure and a full night of sleep occurred after L-DOPA (/ placebo). Therefore, we were able to isolate the effects of dopamine on re-exposure, consolidation and sleep-dependent processing from its effects on initial encoding and retrieval. Items presented only once (Lists *ii, iii, iv*) were expected to have induced weaker memory traces than the re-exposed items (List *i*).

We used d’ (D-prime) as a measure of recognition memory accuracy for each list. D’ is a sensitivity index that takes into account both the accurately detected signal (hits) and inaccurately identified noise (false alarms) (*38*). In other words, d’ captures not just correctly identified “old” words during the recognition test, but it also accounts for incorrect judgements of “new” items as “old”. D’ can be calculated as the difference between the Z-transformed rates of correct hit responses and incorrect false alarms. A higher d’ therefore indicates better ability at performing the task, while 0 indicates performance at chance.

### L-DOPA accelerates forgetting during sleep

L-DOPA given after learning accelerated forgetting of items presented only once when memory was tested the next day (List *ii*) but not at greater delays (Lists *iii, iv,* **Fig. 2a.**). First, we performed pairwise comparisons between the L-DOPA and placebo conditions for each single-exposure list. Data was missing for one participant from Day 5 test following placebo, and therefore we analysed each list separately, as opposed to using an ANOVA which would require removing the participant’s data from all analyses. These comparisons demonstrated that d’ was reduced on L-DOPA (**d’**_List *ii*_ = 1.249 ± 0.59) compared to placebo (**d’**_List *ii*_ = 1.544, ± 0.65) at Day 1 (paired t(34) = −3.333, p = 0.002, BF_10_ = 16.6, n = 35). By Day 3 there was no difference (**d’**_List *iii*_: L-DOPA = 0.86 ± 0.46; placebo = 0.82 ± 0.63; Wilcoxon’s Z = 338, p = 0.313, BF_01_ = 5.2, n =35; **d’**_List *iv*_: L-DOPA = 0.58 ± 0.58; placebo = 0.59 ± 0.55; t(33) = −0.02, p = 0.982, BF_01_ = 5.4, n = 34). The reduction in List *ii* accuracy remained after correcting for false discovery rate for the three tests (Benjamini-Hochberg corrected p = 0.006) (*39*). Together these findings show that L-DOPA accelerates the speed of forgetting for information over 1 night, but this information would be lost in the longer term even without L-DOPA (**Fig. 2a.**, *SM1*). This suggests that dopamine may play an important part in either selecting memories for storage or initiating forgetting.

**Fig. 2.**
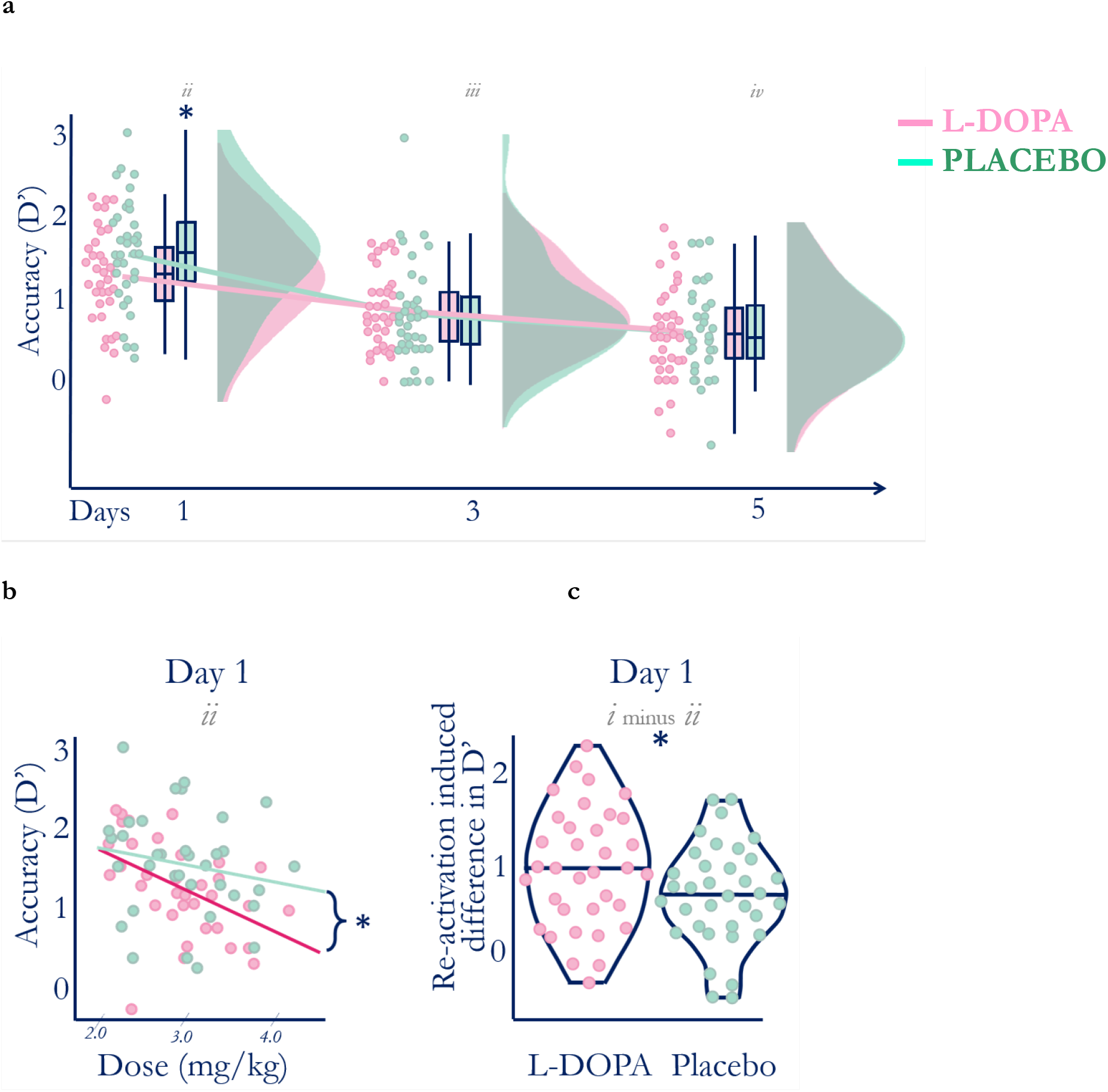
Nocturnal dopamine dose-dependently modulates memory. **a**. Higher d’ at Day 1 on placebo (green) compared with L-DOPA (red) shows that overnight L-DOPA increased forgetting when memory was tested next day (List *ii*) but not 3 or 5 days later (Lists *iii* and *iv* respectively) compared to placebo. Note that L-DOPA was no longer active during memory tests. Therefore, L-DOPA during sleep accelerates forgetting of weakly encoded information that is naturally forgotten by day 3. Boxplot shows quartiles with kernel densities plotted to the right. **b**. Higher L-DOPA dose during consolidation was correlated with poorer Day 1 recall of List *ii* d’ (Spearman’s ϱ = -.056, p < 0.001, red) but no such relationship was found on the placebo night (ϱ = −0.23, p = 0.180, green). Notably, these two relationships were also different (Pearson’s r-to-z transform z= −2.634, p = 0.008) suggesting the effect is driven by dose and not body weight alone. Lines of best fit are shown for illustration purposes. **c**. L-DOPA increased the relative benefit of re-exposed compared to other items (List *i* d’ minus List *ii* d’). This relative benefit was larger after L-DOPA (d’ _List *i–-ii*_ = 0 .953 ± = 0.67) compared to placebo (d’ _List *i–-ii*_ = 0.643 ± 0.56; t (34) = 2.48, p = 0.018, BF_10_ = 2.6). This difference was driven both by an increase in List *i* d’ and a decrease in List *ii* d’ on L-DOPA, although the former was not significant (*SM 3*). Lines show maximum, median, and minimum values (horizontally) and kernel densities (vertically).

We were also interested in any dose-dependent effects of L-DOPA on memory. Body weight is known to influence the cumulative dose and pharmacokinetic properties of L-DOPA (*40*), as well as L-DOPA’s effect on memory in humans (*9*). We used a mixed linear model to investigate the effect of dose (based on body weight). A model with weight-adjusted dose (mg/kg), delay from learning (days) and the interaction term (delay * dose) as fixed effects and participants as random effects (including random intercepts and slopes of delay by participant) revealed a main effect of delay (t(33.7) = −9.142, p < 0.001), no overall effect of dose (t(20.3) = −1.36, p = 0.188) and a delay * dose interaction (t(98.2) = 2.33, p = 0.022, n = 35). The two effects remained following false discovery rate correction for the whole model (*SM 2*). Next, we performed a series of post-hoc correlational analyses to determine which effects were driving this interaction.

The degree of forgetting correlated with L-DOPA dose (Spearman’s ϱ = −0.56, p < 0.001, n = 35, p_corrected_ < 0.001 after correcting for all post hoc correlations) but not with placebo (**Fig. 2b.**– Spearman’s ϱ = −0.23, p = 0.18, n = 35). The degree of forgetting did not correlate with L-DOPA dose (p > 0.36) on days 3 or 5 in either condition. The lack of correlation in the placebo arm suggests that these effects were not driven by bodyweight but were instead associated with the treatment. The delay*dose interaction was therefore driven by L-DOPA affecting memory for List *ii* on Day 1 but not at subsequent delays. This suggests that L-DOPA accelerates initial forgetting in a dose-dependent manner, but it does not influence memory for more strongly encoded items that would be retained 3 or 5 days later.

### L-DOPA accelerates forgetting of weak but not strong engrams

Next, we investigated whether dopamine modulates how re-exposure affects memory. As expected, strong memory traces (re-exposed items – List *i*) were better retained (more ‘hits’) than others (List *ii*) both following L-DOPA (**Hits**_List *i*_ = 18.1 ± 2.1; **Hits**_List *ii*_ = 13.8 ± 3.3; t(34) = 8.49, p < 0.001) and following placebo (**Hits**_List *i*_ = 18.0 ± 2.4; **Hits**_List *ii*_ = 15.0 ± 3.0; t(34) = 7.18, p < 0.001).

While L-DOPA accelerated baseline forgetting for weaker items as shown above (d’ _List *ii*_ = 1.25 ± 0.59) compared to placebo (d’ _List *ii*_ = 1.54 ± 0.11, t(34) = −3.333, p = 0.002, BF_01_ = 0.1, *SM 3*), re-exposed List *i* items were not affected by the treatment (**d’**_List *i*_ = 2.20 ± 0.78; placebo **d’**_List *i*_ = 2.19 ± 0.77; t(34) = 0.134, p = 0.894, BF_10_ = 5.5, **Fig. 2c.**, *SM 4, 5*). Note that we expected that L-DOPA would have enhanced retention for strong engrams while leaving weaker memories unaffected. Instead, L-DOPA accelerated forgetting of weaker information leaving stronger memories largely unaffected.

To quantify this relative effect of dopamine on single-exposed compared to re-exposed items, we used the paired difference between the strongly and weakly encoded lists (i.e. d’ for List *i* minus d’ for List *ii*) from the Day 1 recognition test. This relative benefit was larger following L-DOPA (**d’**_List *i-ii*_ = 0.953 ± 0.67) compared to placebo (**d’**_List *i-ii*_ = 0.643 ± 0.56, (t(34) = 2.48, p = 0.018, BF_10_ = 2.6, **Fig. 2.c.**). Therefore, L-DOPA selectively biased memory retention away from non-repeated items resulting in a larger difference between the two lists on L-DOPA compared to placebo.

To reiterate, L-DOPA differentially modulated strong and weak memory traces, augmenting differences between them. Furthermore, we performed two post-hoc analyses that showed that the treatment had no effect on the false alarm rate (t(34) = 0.527, p = 0.601, BF_01_ = 4.8). Rather, L-DOPA reduced the hit rate (List *ii* – t(34) = −2.89, p = 0.007, BF_10_ = 6.0) – the hits rather than the false alarms drive all the effects of L-DOPA on d’ we identified. This implies that effects of dopamine are related to engram strength rather than modulation of noise that generates false responses.

Importantly, there was no difference in performance during the evening re-exposure tests between placebo and L-DOPA conditions (Day 0 List *i* paired t(34) = .83, p = 0.412, BF_01_ = 4.0, *SM 4*). The Bayes Factor (BF_01_) suggested that these results were 4 times more likely to have been recorded under the null than the alternative distribution. Therefore, dopamine did not affect memory performance before sleep – the effects we report here only manifest after or during the re-exposure.

Together, these findings provide evidence that dopamine biases selection of memories for long term storage by accelerating forgetting of weakly-encoded information, a process that could free memory capacity for storage of more strongly encoded items. Dopamine may bias memory in this way during sleep. Next, we explored polysomnography measures for potential neurophysiological mechanisms underlying dopamine’s effects on memory.

### L-OPA prolongs slow wave sleep

Nocturnal L-DOPA increased time spent in slow wave sleep (stage N3) by ~10.6% (**Fig. 3.a.**) but did not markedly affect the time in other sleep stages or total sleep time (*SM 6*). As most slow wave sleep occurs in the first 4 hours of sleep and the absorption profile of L-DOPA controlled release strongly predicts that dopamine would be increased in the first half of the night (*37*), we expected that dopamine would predominantly affect sleep during this time. As predicted, the observed increase in slow wave sleep occurred only during the first half of the night (as defined by the mid-point between lights-off and lights-on times) on L-DOPA (90.2 ± 34.1 min) compared to placebo (76.8 ± 30.3 min, (t(30) = −3.07, p = 0.005, BF_10_ = 8.7, n = 31, for missing data see *SM 7*). L-DOPA did not affect slow wave sleep duration during the second half of the night (t(30) = −0.387, p = 0.703, BF_01_ = 4.9).

**Fig. 3.**
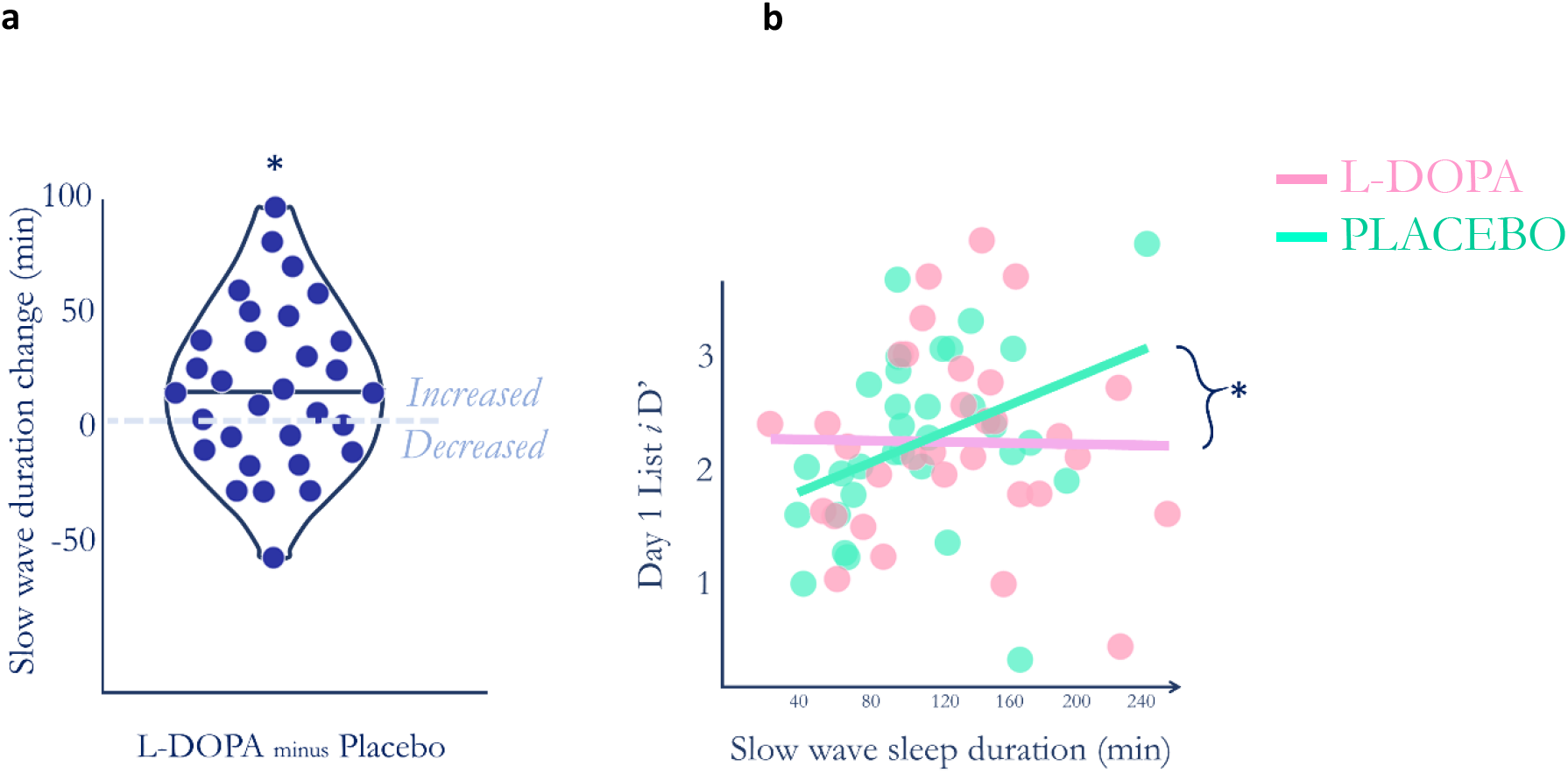
L-DOPA and slow wave sleep duration. **a**. Paired differences in slow wave sleep duration show that most volunteers (dots above zero) had increased slow wave sleep on L-DOPA compared to placebo. The duration was increased by an average of ~10.6% on L-DOPA compared to placebo (t(31) = 2.702, p = 0.011, BF10 = 4.0). This effect remained after false discovery rate correction accounting for each sleep stage (p corrected = 0.044). **b**. Longer slow wave sleep duration was correlated with better memory for strongly encoded information on placebo (Spearman’s ϱ = 0.45, p = 0.009, p _corrected_ = 0.012, green), but after L-DOPA was given this effect disappeared (ϱ = 0.043, p = 0.810, red). The two relationships were different (Pearson’s r-to-z = −1.99, p = 0.046). This strongly suggests that L-DOPA does not increase the relative effect of re-exposure by merely increasing slow wave sleep. Lines of best fit are presented for illustration.

Next, we explored the relationship between L-DOPA’s effect on total slow wave sleep duration with its effects on memory. Overall, longer slow wave sleep duration was associated with enhanced accuracy for the repeated items (List *i*) on placebo (Spearman’s ϱ = 0.450, p = 0.009). This effect did not occur for List *ii* (non-repeated items), and it disappeared after participants took L-DOPA (List *ii* Spearman’s ϱ = −0.043, p = 0.810, **Fig. 3.b.,***SM 8*). This suggests that slow wave sleep duration is associated with consolidation of stronger memory traces, but it does not suggest a direct relationship between slow wave sleep duration and acceleration of forgetting of weaker memory traces on dopamine.

We next asked what neurophysiological processes underlie the quicker forgetting of weaker compared to stronger memory traces on L-DOPA.

### L-DOPA increases spindle amplitude during slow wave sleep

Spindles are a prominent feature of NREM sleep and are associated with memory retention (*41, 42*). L-DOPA increased slow wave sleep spindle amplitude, and while this increase was small on average (m _placebo_ = 28.3 ± 8.5 μV; m _L-DOPA_ = 28.9 ± 83 μV; Wilcoxon’s Z = 95.0, p_corrected_ = 0.008, BF_10_ = 3.6), this effect was manifest in 25 out of 31 participants with spindle data available (**Fig. 4.a.,***SM 9*). This change was not correlated with the weight adjusted dose (Pearson’s r = −0.139, p = 0.456), nor did we find any correlations between spindle amplitude and the d’ difference between Lists *i* and *ii* on either L-DOPA (Spearman’s ϱ = 0.047, p = 0.801) or Placebo (Spearman’s ϱ = −0.040, p = 0.833, *SM 8*). However, the paired change of slow wave spindle amplitude and the d’ difference for strong and weak memories between the L-DOPA and placebo nights was positively correlated (ϱ = 0.438, p = 0.015, n = 30, **Fig. 4.b.**).

**Fig. 4.**
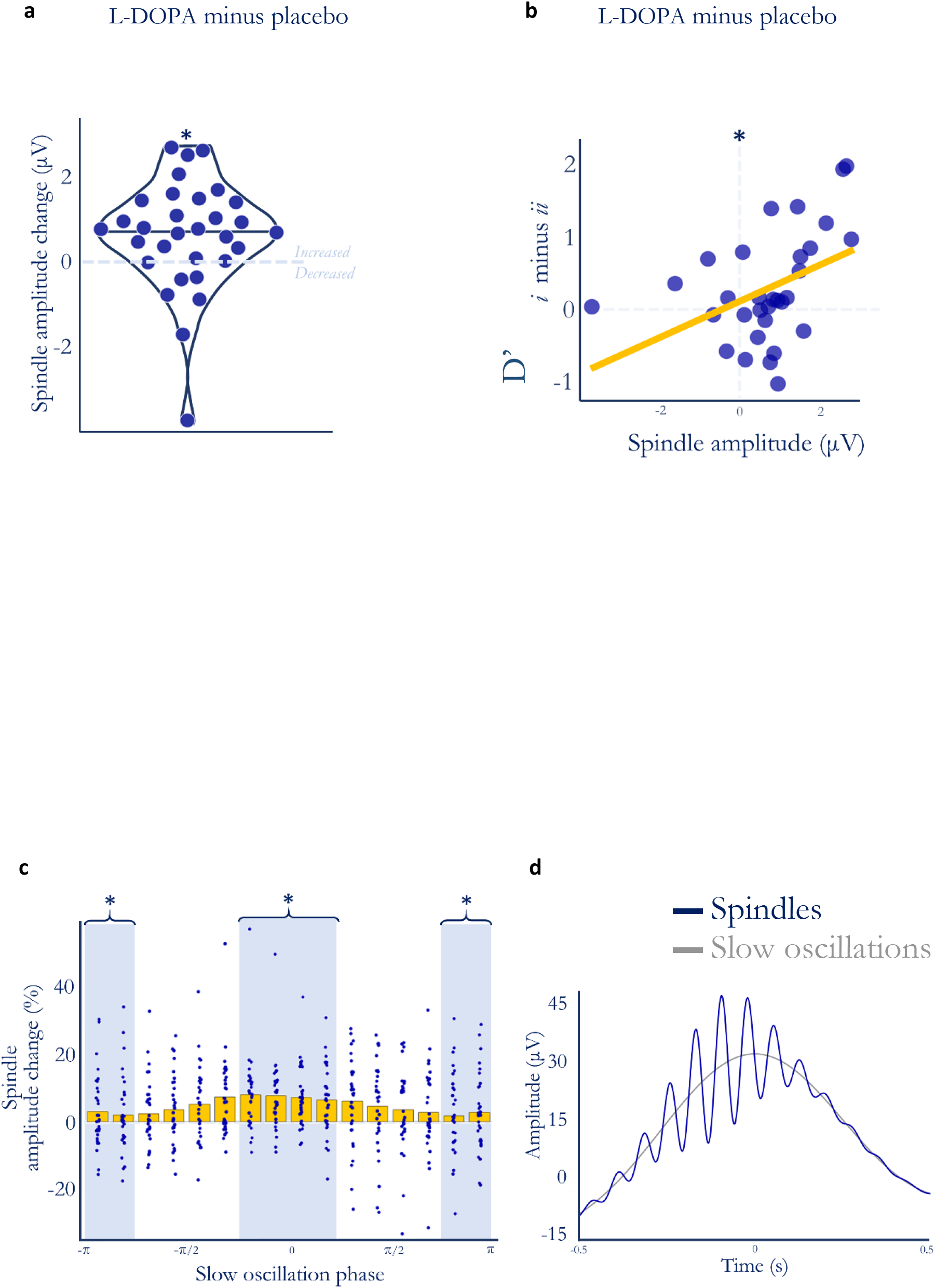
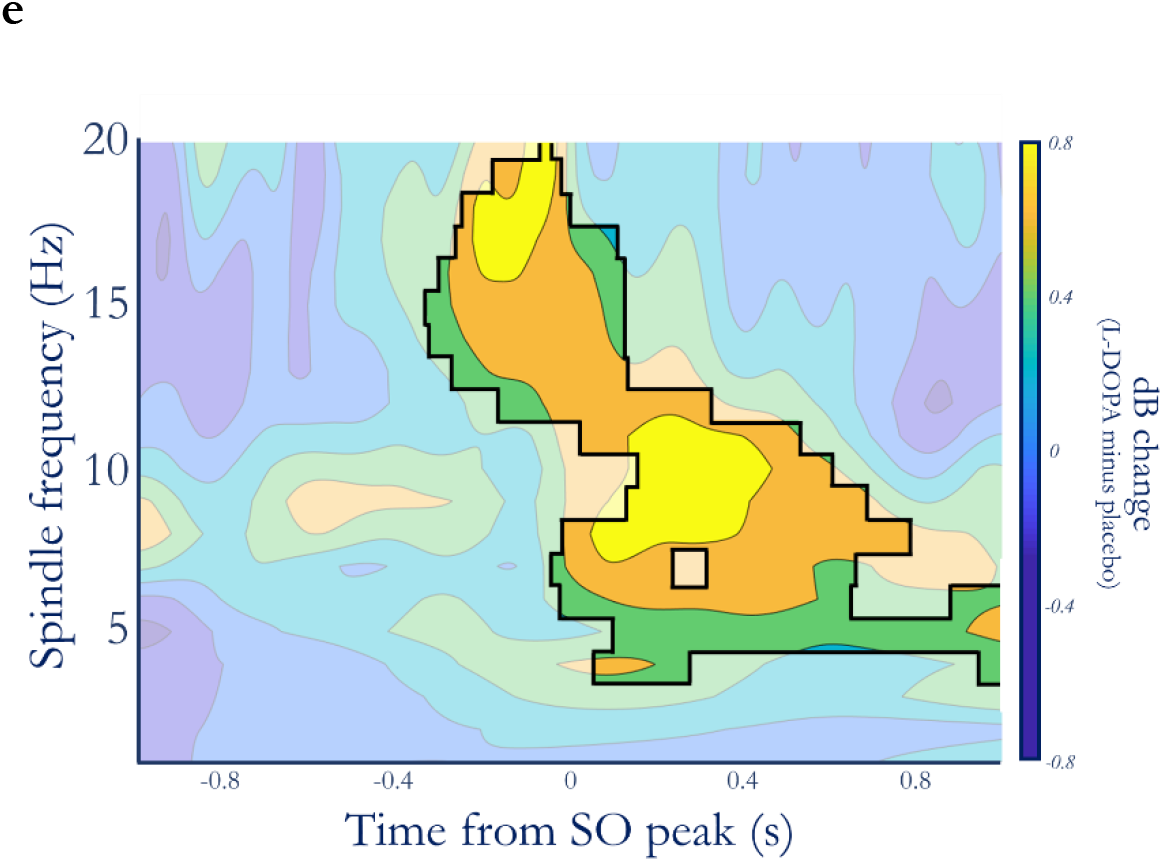
L-DOPA, memory, and spindle amplitude. **a**. Nocturnal L-DOPA increased spindle amplitude (n = 31, Wilcoxon’s z = 401, p = 0.002, p _corrected_ = 0.008, BF_10_ = 3.6) suggesting an effect of L-DOPA on regional coherence during slow wave spindles. **b**. The L-DOPA mediated increase in spindle amplitude was associated with the L-DOPA mediated increase in the relative benefit of re-exposure on d’ (**Fig. 2c.**) (Spearman’s ϱ = 0.438, p = 0.015). Note that this relationship is non-linear, line is fitted in the figure for illustration. **c**. The dopamine-induced spindle amplitude increase is slow oscillation phase-dependent. Mean spindle amplitude change (normalised to baseline amplitude ([placebo + L-DOPA]/2) is higher on L-DOPA around the zero phase of slow oscillations. We compared the effect of L-DOPA at the peak (zero phase) and trough (π phase) of the slow oscillation. The L-DOPA mediated spindle amplitude increase was larger in the 4 zero-centric bins compared to the 4 π-centric bins (outermost on either side) – (paired t(30) = 2.12, p=0.043, BF_10_ = 1.3). Yellow bars show the mean amplitude change with individual participants’ spindle amplitude change overlaid. Spindle amplitude peaked in the -π/4 to - π/8 phase bin for both placebo and L-DOPA. **d**. Peak-locked grand average mean slow oscillation events (grey) superimposed with the peak-locked average of all spindle events (blue) that occurred during slow oscillations – averaged across both L-DOPA and placebo nights. **e**. Paired permutation cluster analysis in time-frequency space comparing L-DOPA and placebo conditions for all slow oscillation - spindle co-occurrence events, centered on the slow oscillation peak. All shown differences denote increased activity on L-DOPA (cluster threshold of α = 0.01, time-frequency space outside significant clusters is greyed). Overall p = 0.002 for the largest cluster.

In other words, the behavioural effect of L-DOPA on memory based on engram strength was associated with the increase in spindle amplitude on L-DOPA. This effect was specific to the L-DOPA-mediated *difference* in memory and spindle amplitude between strong and weak memory traces, and it was not present for List *i* or *ii* alone (*SM 8*).

### L-DOPA affects spindles most at slow oscillations peaks

Temporal coupling between slow oscillations and spindles have been shown to predict memory performance, and this coupling is impaired by aging (*43*). As we observed effects of L-DOPA on both memory and spindle physiology, we next explored whether L-DOPA’s effects on memory performance could be due to an alteration in slow oscillation – spindle coupling.

Periods of time during which the maximum spindle amplitude occurred during a slow oscillation events were identified, segmenting the slow oscillations into 16 phase bins (then grouped into 4 shaded and white areas in **Fig. 4c.).** L-DOPA had a slow oscillation phase dependent effect on spindle amplitude, with a larger increase around the zero phase (**Fig. 4c.**). The peak change occurred in the -π⁄4 to +π⁄4 grouping, the same that showed the highest mean spindle amplitude for both L-DOPA and placebo conditions (**Fig. 4c., d.**).

To see whether this effect was specific to this spindle region-of-interest (ROI)we also performed spectral composition analyses of each slow oscillation – spindle co-occurrence. A spectral mean was calculated for each participant for L-DOPA and placebo conditions using Morlet Wavelet convolution (**Fig. 4e**). Changes in power between L-DOPA and placebo were then identified using a cluster-based permutation method. Cluster analysis of the *a priori* spindle ROI revealed an increase in power on L-DOPA compared to placebo (p = 0.002). **Fig. 4e.** demonstrates the primary cluster (threshold α = 0.01 p = 0.002) when this same analysis was carried out on the surrounding time-frequency space. It suggests that the power increase extends beyond the spindle ROI, into theta and alpha bands following the slow oscillation peak.

L-DOPA therefore altered the neural dynamics that underlie the synchronised relationship between slow oscillations, spindles, and potentially other frequencies. This may represent either a phase-specific effect of dopamine during sleep, or a secondary effect on these dynamics caused by a dopaminergic bias of early awake consolidation or re-exposure.

We found no associations between L-DOPA and other slow oscillation characteristics (all ps > 0.09, *SM 9*). Exploratory analyses revealed no differences between L-DOPA and placebo on subjective sleep measures (St Mary’s Hospital Sleep Questionnaire (*44*) or Leeds Sleep Evaluation Questionnaire (*45*) (*SM 10)*.

### L-DOPA does not modulate memory at encoding or retrieval – Secondary experiment

Whilst there were no differences in performance between L-DOPA and placebo during the re-exposure test in the evening (*SM 4)*, it is possible that the observed effect of L-DOPA on memory was driven by either L-DOPA enhancing encoding during the re-exposure or by residual amounts of L-DOPA acting during retrieval. To investigate whether dopamine was affecting these stages of memory, we ran a secondary placebo-controlled experiment manipulating dopamine levels at encoding or retrieval.

In the secondary experiment, healthy elderly participants were given short-acting L-DOPA an hour before encoding and, in a separate memory task, an hour before retrieval (*SM 11, 12)*. Note that we used short-acting Levodopa here rather than controlled-release (as in the main experiment) as the processes we were probing were discrete events rather than evolving processes. We did not find an effect of L-DOPA on encoding (t(28) = −0.352, p = 0.728, BF_01_= 4.6, n = 32) or retrieval (t(27) = −0.393, p = 0.698, BF_01_= 4.6, n = 28) with a 24-hour delay between learning and test (*SM 12, 14*, for missing data see *SM 7)*. Therefore, at the doses and timings used here, dopamine appears to have a temporally and functionally specific effect biasing memory towards important information after initial learning, during either re-exposure, sleep or both.

Whilst the results from this control study support our initial interpretation that L-DOPA affects memory *after* initial learning but *before* retrieval, it is important to note that due to practical reasons direct statistical comparisons between the two studies cannot be made. First, due to the different profiles of the treatments used (controlled release versus short acting), the L-DOPA doses participants were exposed to were different; 2.1 mg/kg for the control study compared to 2.9 mg/kg for the main experiment. Second, whilst the memory tasks used were similar word-list tasks, they were not identical.

## Discussion

Dopamine accelerates forgetting for weakly encoded information during sleep – while more strongly encoded information is relatively preserved – and increases duration of slow wave sleep by 10.6%. The behavioural effect of dopamine on strongly versus weakly encoded information is associated with a dopamine-driven increase in spindle amplitude during slow wave sleep. This increase in spindle amplitude only occurs around the peak of slow oscillations.

Traditionally, forgetting is considered a passive process where information is “lost”. However, newer animal models support an active, more strategic, forgetting process mediated by dopamine (*31, 33, 34, 46*). We showed that dopamine increased forgetting for information at 1-day delay but not at later timepoints. Therefore, dopamine may accelerate forgetting of low importance information that would inevitably be lost over time. Our data suggest an *active* dopamine-dependent forgetting mechanism in humans – which can be conceived as dopamine biasing memory selection away from weakly encoded items. This may in turn allow prioritisation of strongly encoded or salient items for consolidation.

Such prioritisation may be further explained – through analogy with drosophila experiments – by a second dopaminergic system that protects important information from forgetting (*46*). Human behavioural evidence supports preferential consolidation of salient or rewarded information during sleep (*19, 47, 48*), and we tie this more closely to dopaminergic modulation. Contrary to hypothesis,, we did not find evidence for a dopamine-driven direct enhancement in consolidation of strongly encoded information here. The relative effect of dopamine on forgetting of low versus high importance items suggests a more nuanced dopaminergic effect - biasing memory away from weaker memory traces Given L-DOPA is already widely used in clinical practice for Parkinson’s disease and could be quickly repurposed if this effect is beneficial for memory overall, this certainly warrants attention and further clinical research.

There is clear evidence that memory processes before sleep can alter slow wave sleep characteristics, particularly in the early part of the night (*49*). We administered dopamine while participants were awake, 2h before their bedtime, thus it is possible that at least a portion of the dopamine-driven increase in forgetting occurred during wake. Whilst we cannot rule this out, we did observe that L-DOPA compared to placebo was associated with changes in scalp electrophysiology during sleep, with some of these effects being associated with memory performance. Therefore, we suggest that the dopamine-driven changes on memory were sleep-dependent.

While changes in spindle characteristics are well known to be associated with memory and neurodegeneration (*50*), this study directly links dopamine with behavioural relevance of spindles. Spindle amplitude is shaped by the interplay between the thalamus and the cortex (*51*), and increased amplitude reflects a more coherent and wider topographical expression of spindle-related activity, i.e. better coordination between the brain regions (*52, 53*). Behaviourally, spindle amplitude has also been associated with enhanced memory retention during a motivated forgetting task (*54*) and during a tagging paradigm (*55*). This coordinated activity between the thalamus and cortex during sleep may thus be associated with selecting memories for later retention. Consistent with this, we showed that greater spindle amplitude was associated with a larger dopamine-induced difference between retention of strongly and weakly encoded information.

L-DOPA mainly increased spindle amplitude around the peak of slow oscillations, which occurred despite no change in slow-oscillation amplitude. Spindles peaked just before zero phase, consistent with previous findings in healthy elderly (*43*). Spindles, particularly when nested in slow oscillation peaks, are hallmarks of sleep-dependent memory consolidation (*56*). Age-related uncoupling of spindles from peak of slow oscillations increases overnight forgetting (*43*). We interpret dopaminergic increase in spindles synchronised to near zero phase of slow oscillation as enhancement of physiological spindle activity to modulate memory consolidation.

There are two possible explanations for our finding – (1) dopamine directly enhances spindle amplitude which in turn enhances the way in which memory is biased based on encoding strength (2) or dopamine during memory re-exposure before sleep results in stronger behavioural tags that in turn alter subsequent spindle amplitude to reflect the changes in the memory engram that took place during tagging. These effects are not mutually exclusive, and indeed could be interacting. Future experiments separating the effects of sleep consolidation from re-exposure benefit are necessary to disentangle this.

We suggest that two simultaneous processes may be at play (**Fig. 5.**). First, during learning a portion of information is “tagged” as important (*57*), and dopamine enhances this process by creating a stronger tag (*58, 59*). Second, during subsequent sleep, dopamine increases forgetting for the less important, non-tagged items while the tag shields the important (or re-exposed) information from forgetting (*60, 61*). This theory has been proposed before, and the current study adds to it by implicating (dopamine-mediated) crosstalk between the thalamus and the cortex during spindles as a potential mechanism for the later effects.

**Fig. 5.**
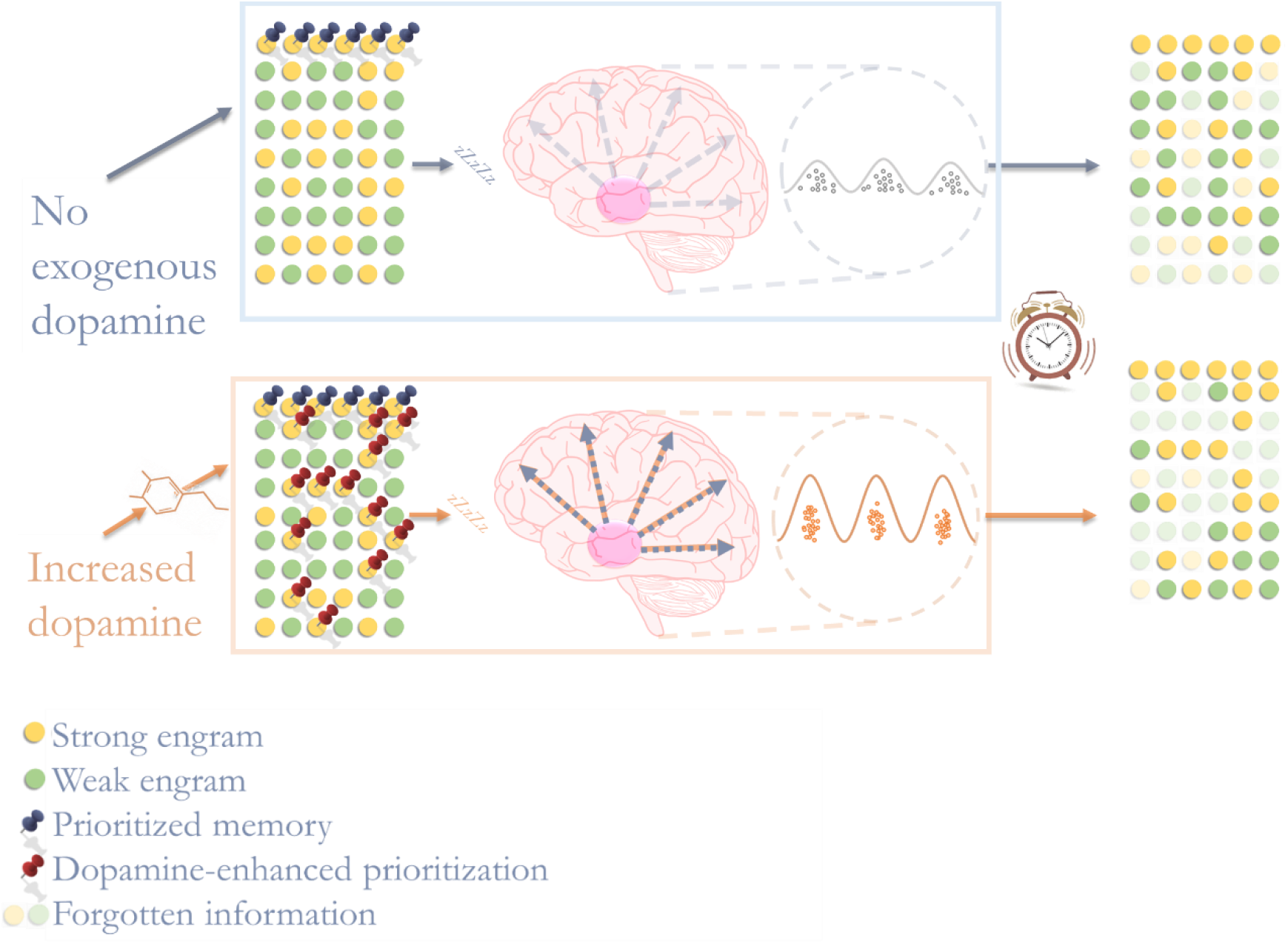
Dopamine modulates memory after learning by enhancing memory prioritisation and subsequent sleep processes. A proportion of important information (yellow engram) is earmarked for retention by a neural “tag” during re-exposure (blue pin). Dopamine during re-exposure enhances this effect (red pins) at the expense of unpinned engrams (green). During sleep, weak engrams are preferentially forgotten whilst ear-marked information remains unaffected, leading to a more selective memory trace. Dopamine modulates these selective memory processes by enhancing synchronisation in cortical firing patterns during spindles, at the peak of slow oscillations. Together these two processes (enhanced prioritisation and synchronisation) bias subsequent memory.

Our observed findings may be specific to older people. Memory loss is a prominent problem in old age and our eventual goal is to improve quality of life through cognitive enhancement, justifying the use of a target population of interest to future trials. There is drop-out of dopaminergic neurons that comes with old age (*62–64*) (*62–64*) which has been shown to affect the impact of taking dopaminergic medications on cognition (*65*). Ageing decreases the duration of slow wave sleep, and the number and amplitude of spindles (*66*), with some reporting nearly a 50% reduction in spindle amplitude with advanced age (*67*). Models in non-aged animals suggest that D2 receptors promote wakefulness (*68*) and dopamine levels are generally higher during wake than sleep in animals (*69*). In young healthy adults, direct administration of a dopamine *antagonist* during slow wave sleep actually increases the duration of slow waves sleep (*70*). It has been noted before that the wake-promoting effects of dopamine in the young contradict the sleepiness that is a recognised side effect of L-DOPA in patients with Parkinson’s disease (*71*). Therefore, age should be considered when interpreting the effects of dopamine on memory and sleep.

Furthermore, slow wave sleep may be affected early in Alzheimer’s Disease (*72*). Interrupting slow wave sleep is proposed to hinder clearance of amyloid from the brain and amyloid plaques are one of the key pathological changes in Alzheimer’s Disease (*73, 74*). L-DOPA is routinely prescribed for Parkinson’s disease with a good safety profile; however, the impacts of L-DOPA on sleep have not been assessed in detail except in small studies of Parkinson’s disease (*75–77*). Our finding that L-DOPA may ameliorate age-dependent spindle loss with concomitant memory benefits could be promising for treating age-related memory decline, or more severe memory deficits found in Alzheimer’s dementia. Perhaps more excitingly, our current findings may have implications for prevention of Alzheimer’s disease. Through increasing slow wave sleep duration and spindle amplitude with nocturnal dopamine, we open up a new therapeutic avenue for Alzheimer’s disease prevention – repurposing L-DOPA to prevent Alzheimer’s.

Together, our findings suggest that the repetition-benefit on memory is improved by dopamine at the time of the repetition and during sleep-consolidation, which is mediated by increased slow wave sleep duration and spindle amplitude. We propose that this dopamine-induced increase in spindle amplitude reflects more synchronous cortical activity during spindles increasing forgetting of weakly encoded items and with a net effect of augmenting the difference between strongly and weakly encoded engrams (**Fig. 5.**). These findings have potential clinical impact in enhancing sleep and memory selection in old age, and in mild amnesic disease.

## Method

### Participants

We recruited 70 elderly (65+ years) volunteers to complete the two studies reported here (n = 35 each study, see *SM 4, 15, 16*). All aspects of this research adhered with the Declaration of Helsinki and we had relevant ethical and regulatory (UK) approvals in place (Study 1 ISRCTN: 90897064).

### Design

#### Study 1

In the main placebo-controlled double-blind randomised study, volunteers were initially screened over the phone for common exclusions, and then invited for three in-house visits. On the first visit they were fully screened for eligibility, and they practiced the memory task. They were asked about their usual sleeping pattern so that the second and third visits could be designed to follow each participants’ usual sleep routines as much as possible. On the second visit, volunteers arrived on site in the evening where they were re-consented and screened for continued eligibility. For an outline of the evening see **Fig. 1a.**, *SM 11*.

First, volunteers learnt a verbal memory task (**Fig. 1b.**). Thirty minutes after learning, they were given 200mg L-DOPA or placebo. 75 min after dosing, a quarter of the items (List *i*) were re-exposed by a recognition test where no feedback was given. The purpose of this test was to create a stronger memory trace. 45 min after the re-exposure the volunteers went to bed. Each evening was designed based on each participants’ usual sleeping pattern (L-DOPA administered 2h prior to switching the lights off for the night at their usual bedtime).

Volunteers slept on-site for a full night, and they were woken up at their usual wake-up time. Around 1.5h after waking up, approximately 12h after dosing, volunteers’ verbal memory was tested again (Lists *i* and *ii)* before they left the study site. 2 and 4 days later (3 and 5 days after learning) they were contacted over the phone for follow-up recognition memory tests (for Lists *iii* and *iv*, respectively).

The second and third visits were identical except for treatment (L-DOPA / placebo) allocation. This study obtained ethical approval from the South West Central Bristol NHS Research Ethics Committee (REF: 16/SW0028) and clinical trial authorisation from the Medicines and Healthcare products Regulatory Agency (IRAS ID:178711).

#### Study 2

In the secondary placebo-controlled double-blind crossover experiment, volunteers were first screened over the phone before inviting them on site for the test sets. Each test set carried over for three days: On the Day −1 (relative to dosing) participants learnt word-list on site, on Day 0 they were dosed with 150mg L-DOPA (or placebo, *SM 11, 12*) and then tested on the previously learnt words (retrieval test). Immediately following this, they learnt another word-list (encoding condition) for which their memory was tested the following day (Day 1) over the phone. Therefore, participants learnt two word lists on each test There tests were timed so there was approximately 24h in between learning and test.

This study obtained ethical approval from the University of Bristol Faculty of Medicine and Dentistry Ethics Committee (REF: 12161).

### Treatment

#### Study 1

In the main placebo-controlled randomised double-blind study, each participant was dosed with co-beneldopa controlled release (containing 200mg L-DOPA) was given in capsule form and placebo (encapsulated inert powder, matched for appearance). Blinding and randomisation were performed in blocks of 6 by author LM, Production Pharmacy, Bristol Royal Infirmary, University Hospitals Bristol and Weston NHS Trust. On the study nights, dose was given by an on-site medic who was blind to treatment condition and played no role in collecting data. The treatments were given at different visits. Both treatments were preceded by Domperidone 10mg (tablet) to alleviate possible nausea caused by L-DOPA.

#### Study 2

In the L-DOPA condition of the secondary study volunteers received 10mg of Domperidone (anti-emetic) 30 minutes before co-beneldopa (containing 150mg L-DOPA). Both medications were dispersible, and they mixed into cordial to hide taste and residue. In the placebo condition, volunteers received plain cordial in place of Domperidone and vitamin C mixed into cordial in place of L-DOPA. Blinding and randomisation were performed by members of the research group who had no other involvement in this study.

While the two experiments were designed to complement one another, for practical reasons there were several important differences in study designs. First, the L-DOPA given in the main study was long-acting and of higher dose (4-8 hours cf. 1-4 hours and 200mg cf. 150mg) to target consolidation during sleep which is a longer process than encoding or retrieval. Second, the controlled release L-DOPA in the main study was encapsulated, whilst in the secondary study we used dispersible L-DOPA. For this reason, the placebo used in the main experiment was encapsulated inert powder, whilst in the second study we used dispersible vitamin C. These differences and individual differences in dopamine absorption and metabolism introduce unmeasurable differences between the two experiments that need to be considered when interpreting differences between them.

### Verbal memory test

#### Study 1

Volunteers learnt four lists (*i, ii, iii,* and *iv*) of 20 target words (total 80 targets) presented on a computer screen one at a time, in a random, interleaved order (**Fig. 1b.,***SM 12*). Each word was presented once for 3.6s during which the volunteers were asked to determine if the items were alive or not to assist learning. They were instructed to remember as many of the words as they could.

During test phases, volunteers were presented with a list of 40 (days 0, 3, and 5) or 80 words (day 1), half of which were targets (present at learning) and half of which were distractors (not presented previously). They were asked to judge whether words were targets or not. On days 0, 1, 3, and 5 memory was tested for Lists *i*, *i* and *ii*, *iii*, and *iv* respectively. Therefore, List *i* was tested twice: First in the evening while L-DOPA (/ placebo) was active in the system and then again in the morning together with List *ii*. The re-exposed and novel (List *i* and List *ii*, respectively) targets tested on day 1 were assessed to study L-DOPA’s effect on behavioural tagging of ‘important’ information. The rationale was that when a word is presented a second time (during re-exposure), it will be deemed more important and will be preferentially remembered. The distractors were unique at each test.

#### Study 2

The purpose of this study was to test L-DOPA’s effects on retrieval and encoding. Two separate memory tests were conducted (*SM 13).*

##### Retrieval

During learning on D-1 (day before dosing) volunteers were presented with 48 complete nouns on a computer screen. They were instructed to read the words aloud and try to memorise them for later. Each word was shown once for 5 seconds separated by a fixation cross in the middle of the screen for 2 seconds and no responses to the words were made during learning. There were no breaks in the learning block (total duration = 5mins 36secs). Memory was tested using unique words 30 minutes (D-1, baseline) and 24 hours (D0) after learning. The D0 test was given when L-DOPA was at its peak concentration (~ 1h following dosing). In the test phases (D-1 and D0). This test took approximately 5 minutes to administer.

##### Encoding

D0 around 1.5 hours after dosing, after the test for the previous task had finished, volunteers saw a list of 96 complete nouns presented on the computer screen. Each word was displayed for 5 seconds, followed by a fixation cross for 2 seconds. The words were first presented in a random order over two blocks, and then again in a different random order over two more blocks, each word was presented twice (n blocks = 4, n words per block = 48, n breaks = 3, block duration = 5 min 36 s). Memory was prompted immediately after learning (D0), and 1, 3, and 5 days later. Each target was tested once with unique distractors (*SM 14*).

Across experiments, learning and tests were completed on a laptop on-site, or over the phone. The experiments were programmed in the MATLAB environment (2015b or 2017a) using the Psychophysics Toolbox V3 (*78*). The scripts and data are available from corresponding authors upon request.

### Polysomnography

In the main experiment, standard in-laboratory polysomnography, including video, was recorded during both study nights using the Embla N9000 amplifier and Embla RemLogic software (Natus Medical Inc., California) at CRIC Bristol, University of Bristol, Bristol, UK. We recorded 12 scalp EEG channels (F3, Fz, F4, C3, Cz, C4, M1, Pz, M2, O1, O2, and a ground electrode approximately between Cz/P3 and C3/Pz) placed according to the 10-20-20 system. Eye movements were detected by electro-oculogram recorded from E1 and E2 sites, and muscle tone from electromyogram recorded below the chin. A 2-lead ECG was also recorded. All signals were sampled at 500Hz. The recordings started 2.5h after dosing when lights were switched off for the night and continued until the volunteer woke up.

### Analysis

#### EEG

##### Event scoring

Sleep stages in 30s epochs were identified manually in accordance to standard criteria (*79*) by two expert scorers, and a third scorer visually assessed a random 10% of ratings for quality. Durations of N1, N2, N3 (i.e. slow wave sleep), REM, awake, asleep and total time in bed were extracted in minutes. First and second halves of the nights were defined by the middle time-point between switching lights ON and OFF. When there was an odd number of epochs, they were rounded so that the first half of the night had the extra epoch.

##### Spindle detection

Spindle characteristics were then isolated with in-house written MATLAB scripts using the EEGlab toolbox (*80*). Electrodes were re-referenced to contralateral mastoid and empty and high variance epochs were removed. Thereafter, only data from the Cz electrode was used. First, the channel was visually inspected and epochs with high noise or clear artefacts were removed manually. Data was then filtered (high pass 11Hz, low pass 17Hz) and rectified.

Next data was smoothed using a moving average window of 200ms before down-sampling to 100Hz (from 500Hz) for computational efficiency. An event was marked as a spindle if the threshold exceeded the 90th percentile for that data set (i.e. sorting data into an ascending order and including top 10%) for .5 – 3 seconds, with a minimum of 0.5s between spindle events.

##### Slow oscillations

The slow oscillation detection process followed the same re-referencing and noise removal methods used for spindle detection, without smoothing. Data from the CZ electrode was filtered between 0.16Hz and 1.25Hz and then z-scored. We applied a threshold of 75%; if the slow oscillation amplitude surpassed this threshold for 0.5 - 5 seconds (including multiple events if separated by <0.25s), it was marked as a slow oscillation. The duration of the event was determined by the closest oscillation maxima following the amplitude dropping below a 60% threshold on each side.

##### Spindles and slow oscillations

We identified spindle-slow oscillation co-occurrences as cases where the maximum amplitude of a spindle event coincided with a slow oscillation event, again using the CZ electrode. Using the time stamp of the spindle max amplitude as the centre point, we calculated how spindle amplitude varied with slow oscillation phase over one cycle. First, we divided the oscillation events into 16 bins, equally distributed in phase space around zero, to calculate how the spindle amplitude varied with slow oscillation phase for each coinciding case (for statistical analysis we grouped together 4 adjacent frequency data points (bins) to generate 4 bins as shown in **Fig. 4c**).

The spectral composition of each was done using a Morlet wavelet time-frequency method over a 4s window centred on the slow oscillation peak. Morlet waves at 20 frequencies were used, with 3 cycles for the lowest frequency (1Hz) and 6 cycles otherwise (2-20Hz). A spectral mean was next calculated for each participant for L-DOPA and placebo conditions (**Fig. 4e.**). A cluster-based permutation method (permutation n = 500, cluster threshold of α = 0.01), implemented with the Fieldtrip toolbox (*81*), was used to identify power differences. Based on the finding that max spindle amplitude occurring near zero slow oscillation phase predicts to memory performance in ageing {Helfrich, 2018 #3297}, an a-priori spindle region of interest of 11 - 16Hz, −0.5 - 0.5s was chosen for initial analysis. This same cluster method was then carried out on the wider time-frequency space, 1 – 20Hz, −1 – 1s, the primary cluster (p=0.002) of which is shown in **Fig. 4e.**.

## Behaviour

**Pairwise comparisons** (placebo versus L-DOPA) were calculated using either t-tests or Wilcoxon’s rank tests in R 3.5.3 using RStudio. We also employed a Bayesian paired t-tests in JASP 0.9.2.0 (*82*) to obtain Bayes Factors (BF) – this allows more meaningful estimates of confidence in both significantly different and null results than standard t-tests. BF gives the probability of the data under either hypothesis. E.g. a BF_10_ of 5 would denote that the data is 5 times more likely to have been sampled from the alternative compared to the null distribution, while a BF_01_ of 5 would denote that the data is 5 times less likely to have been sampled from the alternative compared to the null distribution (i.e. 01 versus 10). We defined the prior (expected) distribution as a Cauchy distribution with a mean of 0 and an interquartile range of .5 [δ~ Cauchy (0, .5)]. In other words, we predicted that the δ lies between -.5 and .5 with a 50% confidence. We selected this one as the δs in cognitive neurosciences typically are within those bounds, and as we did not have an informed prediction for the effect sizes.

All mixed modelling was performed on R 3.5.3 using Rstudio, lme4 (*83*) and lmerTest (*84*). We included the participants as random effects and the dose (mg/kg) and the memory test delay (Day 1, Day 3 and Day 5), or memory strength (re-enforced versus not), depending on the analysis, as fixed effects. All fixed effects were mean-centred but not scaled. We selected the model using the maximum feasible fit as this has previously been shown to be the best approach for confirmatory hypothesis testing (*85*).

## List of supplementary materials

SM 1 : Paired differences for delayed memory test

SM 2 : L-DOPA accelerates forgetting.

SM 3 : Re-activated items better retained in both conditions

SM 4 : Pairwise accuracy for List i across tests

SM 5: L-DOPA has disparate effects on forgetting rate depending on whether items were re-exposed or not

SM 6 : Single dose of nocturnal L-DOPA increases time spent in slow wave sleep by 10.6%.

SM 7 : Missing data

SM 8 : Sleep and memory correlations on L-DOPA and placebo

SM 9 : L-DOPA increases spindle amplitude

SM 10 : Subjective sleep measures

SM 11 : Sleep visit timeline

SM 12 : Verbal Memory Task

SM 13 : Secondary study timeline

SM 14 : Study 2 results

SM 15: Demographic information

SM 16 : Exclusion and inclusion criteria

SM 17 : Secondary study results

SM 18 : Individual Power Change

SM 19 : CONSORT diagram

## Acknowledgements

First, we would like to thank each of our participants for their time. We would also like to thank doctors Nicholas O’Donnelly, Lisa Knight, Hilary Archer, Elizabeth Mallam, David Evans and Tom Dalton for providing clinical cover for this study, and doctors Claire Rice and Catherine Pennington for their time and guidance as the steering committee. We are grateful for Alex Howat, Nerea Irigoras Izagirre, and Will Mears for their substantial help in collecting the data. We would like to thank Phil Boreham, cardiologist, for advising us on the safe use of domperidone. We would also like to thank Join Dementia Research, Bristol Brain Centre, North Bristol NHS trust, and CRICBristol staff, and our funders; the Medical Research Council, BRACE Bristol and the David Telling Trust.

## Author contributions

HKI and EJC designed Study 1. MWJ, CD, CO, and LM contributed significantly to designing Study 1. Randomisation and blinding were performed by LM for Study 1. HKI, JPG and EJC designed Study 2. HKI and JPG developed the verbal memory tasks. HKI, GA, JPG, WJC and UB wrote all analysis scripts. Sleep scoring was performed by WJC and OR and overseen by HKI, spindle and slow oscillation analyses were carried out by HKI, WJC and UB. CO and UB gave further statistical guidance. All data collection was overseen by HKI, JPG and EJC. Data was collected by HKI, WJC, GA, OR, JS, RW, EF, JM, CD, ARW, CMN, and JPG. Further clinical cover was provided by EJC, JM and JS. HKI, EJC, WJC, JPG, CO, MWJ and UB interpreted the data. HKI and EJC wrote the manuscript, all authors contributed to the editing of the manuscript and approved of the final version.

## Data availability

Contact the corresponding authors for copies of the MATLAB and R scripts used in analysis, the experimental standard operating procedures, MATLAB scripts for the verbal memory tasks, or word list. Data will be shared in line with sponsor’s requirement for availability of anonymised datasets from clinical trials. The data for the control study can also be shared upon request.

## Code availability

Contact the corresponding authors for copies of the MATLAB and R scripts used in analysis.

