## Supplementary materials for "Dopamine-gated memory selection during slow wave sleep"

SM 1 : Figure - Paired differences for delayed memory test

**Single exposure (List *ii*, List *iii*, List *iv*)**

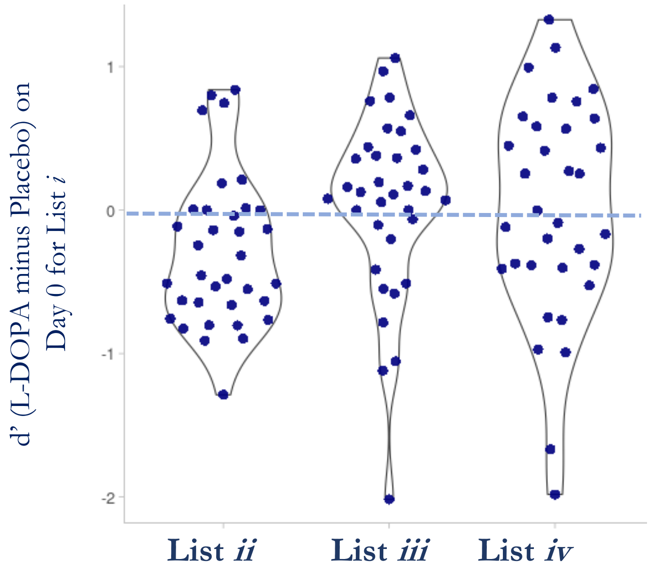

This figure complements **Fig2a** in the main paper. **Fig2a** maps the individual performance for each participant for L-DOPA and placebo nights separately. Here, you can see the paired difference in performance. The relevant inferential statistics are reported in the main text and in Supplementary Table 1. Blue line denotes no difference. Data points above the blue line performed better following L-DOPA compared to placebo.

**Descriptive Statistics**

| | **D1** | **D3** | **D5** |
|---|---|---|---|
| **Mean difference (L-DOPA minus placebo)** | -0.294 | 0.037 | -0.003 |
| **Std. Deviation** | 0.523 | 0.629 | 0.762 |
| **Minimum** | -1.288 | -2.017 | -1.982 |
| **Maximum** | 0.839 | 1.060 | 1.327 |

### SM 2 : Table - L-DOPA accelerates forgetting.

**A**

| | **Estimate** (std error) | **t** | **df** | **p** | **p** (corrected) | **Forgetting $R^2$** marginal |
|---|---|---|---|---|---|---|
| | Accuracy ~ Delay * Dose + (Delay + Dose \| \| participant) | | | | | |
| Intercept | .940 ( .06 ) | 14.931 | 34.0 | <.001 | <.001 | |
| Delay | - .808 ( .09 ) | -9.142 | 33.7 | <.001 | <.001 | |
| Dose | - .031 ( .02 ) | -1.364 | 20.3 | .188 | .188 | .258 |
| Delay * Dose | .123 ( .05 ) | 2.325 | 98.2 | .022 | .029 | |

**B**

| | **Mean** (SD) L-DOPA | Placebo | **Credible interval δ** | **df** | **t** | **p** | **p** (corrected) | **$BF_{01}$** | **$H_0$ vs $H_1$** |
|---|---|---|---|---|---|---|---|---|---|
| Day 1 | **1.249** ( .59 ) | **1.544** ( .65 ) | [ - 1.202 – - .232] | 34 | - 3.333 | .002 | .006 | <.1** | 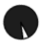 |
| Day 3 | **.855** ( .46 ) | **.818** ( .63) ) | [ - .360 – .508 ] | 34 | 337.5* | .313 | .470 | 5.2 | 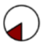 |
| Day 5 | **.584** ( .58 ) | **.593** ( .55 ) | [ - .434 – .428 ] | 33 | - .023 | .982 | .982 | 5.4 | 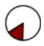 |

**L-DOPA accelerated forgetting when memory was prompted 1 but not 3 or 5 days after learning.**
**A.** Both delay in days and L-DOPA status explained variability in accuracy (d'). $R^2$m quantifies extent of variance in accuracy explained by the fixed effects, their interactions and the intercept. Note that estimates are mean-centred. We opted to use a mixed linear model as we had one missing data point (1 participant, day 5) and more traditional approached (e.g. ANCOVA) would require excluding all data from this participant. Top line of table A denotes model specification in R. **P-values are FDR corrected** (test n = 4) for the whole model using Benjamini-Hochberg procedure.
We included delay (days 1, 3 or 5), dose (mg/kg) and delay * dose interaction as fixed effects, with participants as random effects (including slopes and intercepts) in a mixed linear model. The delay * dose interaction ($p = .022$) explained variability in accuracy, but with no main effect of dose. Post-hoc correlational analyses revealed that higher doses were associated with poorer performance in the L-DOPA ($\varrho = -.56$, $p < .001$) but not in the placebo arm ($\varrho = -.23$, $p = .18$), and that these two relationships were different ($z = -2.634$, $p = .008$, (*1*)). Dose was not associated with performance on other days. As dose was calculated using body weight, it is noteworthy that as it is not associated with measures on placebo, these effects are likely to be drug-related and unlikely to be driven by differences in body size.
**B.** These tests were post-hoc and exploratory. L-DOPA compared to placebo administered after learning lowered d' on day 1 but not at later times. $BF_{01}$ and $H_0$ vs $H_1$ show the probability of our data having been observed under the null (white) as opposed to alternative (black) hypothesis. **P-values are FDR corrected** for days (test n = 3) using Benjamini-Hochberg procedure. Δ denotes effect size for the paired differences derived from the Bayesian posterior distribution. Credible intervals overlapping zero denote no difference. All errors <.001%. Note that $BF_{01}$ which denotes how much likelier our data are under the null are reported as opposed to $BF_{10}$ for easier interpretation.
* Wilcoxon test used due. The sample contained zero values for paired differences. P-values for Wilcoxon tests for such data are less reliable
** $BF_{10}$ = 16.6 (our data is 16.6 times more likely to have been drawn from the alternative than the null distribution)

SM 3 : Figure - Re-activated items better retained in both conditions

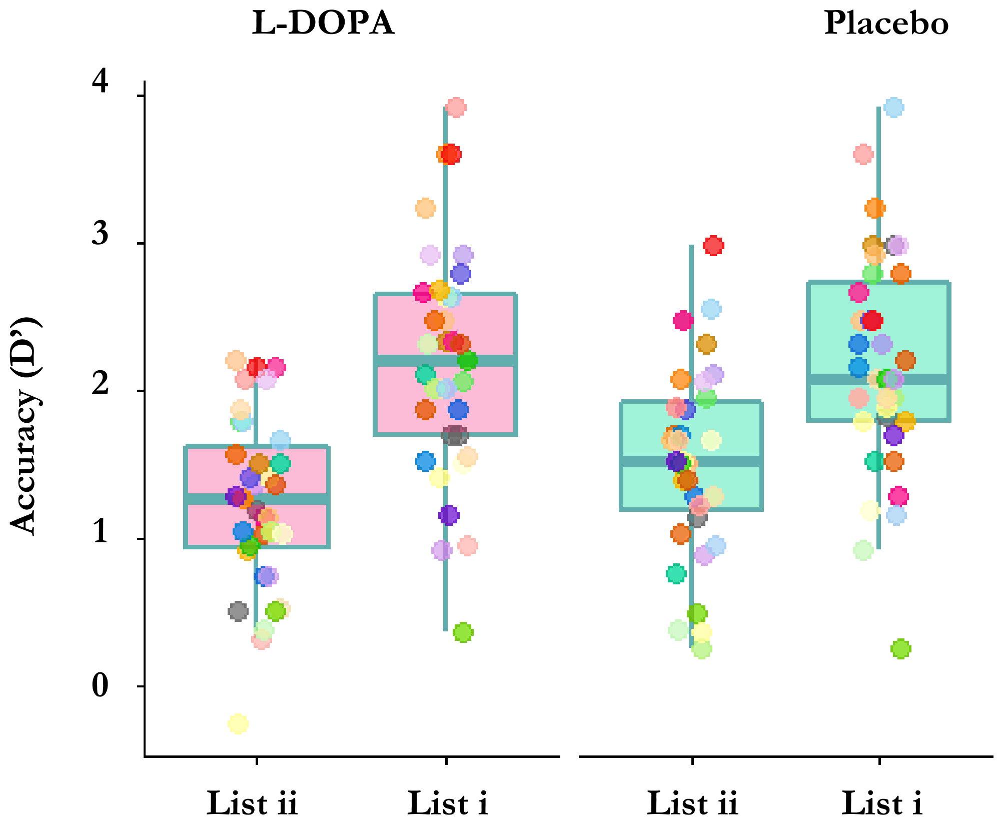

**Re-activated (List i) items better remembered in both conditions**

Performance for re-exposed compared to learnt items (List i to List ii respectively) was better both ON and OFF L-DOPA. On both the L-DOPA (t(34) = -8.419, p<.001, $BF_{10}$ = 14300000, error <.001%) and on placebo (t(34) = -6.764, p<.001, $BF_{10}$ = 165589, error <.001%) re-activated items ($mean_{L\text{-}DOPA}$ = 2.203 ± 0.78, $mean_{placebo}$ = 2.187 ± .77) were better retained on Day 1 than items that were not re-activated ($mean_{L\text{-}DOPA}$ = 1.249 ± 0.59, $mean_{placebo}$ = 1.544 ± .11).
Boxplot lines show median and quartiles. Individual datapoints plotted

### SM 4 : Figure - Pairwise accuracy for List *i* across tests

#### a) Baseline performance on Day 0

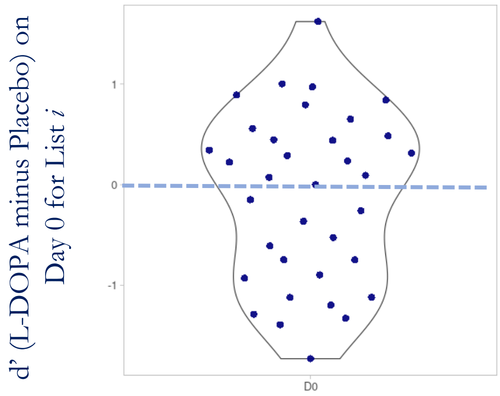

#### b) Memory performance on Day 1

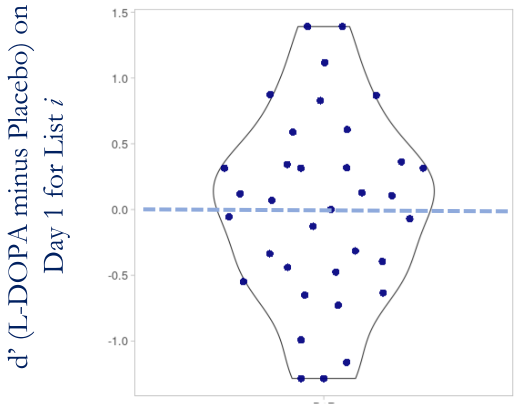

#### c) Performance change from day 0 to day 1 ( day 1 minus day 0)

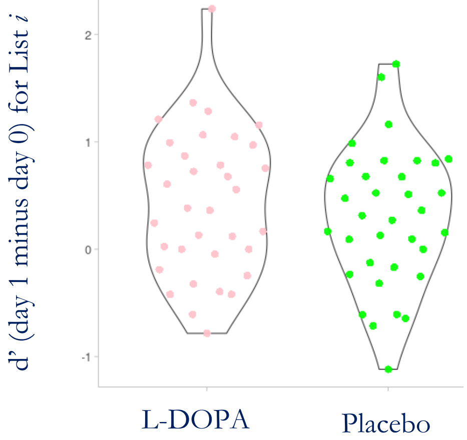

| | Re-exposure | Day 1 test | Change on L-DOPA | Change on placebo |
|---|---|---|---|---|
| **Mean** | -0.118 | 0.016 | 0.432 | 0.298 |
| **Median** | 0.072 | 0.070 | 0.384 | 0.315 |
| **Std. Deviation** | 0.839 | 0.704 | 0.671 | 0.644 |

All of these analyses are exploratory. The table shows summary statistics corresponding to data in figures. All n = 35

a. there was no difference in performance on baseline, t(34) = 0.831, p = 0.412, BF10 = 0.25, BF01 = 4.0, error <0.001
b. there was no difference in performance on day 1, t(34) = 0.134, p = 0.894, BF10 = 0.18, BF01 = 5.47, error <0.001
c. there was no difference in change in performance, t (34) = 0.906, p = 0.371, BF01 = 3.7, BF10 = 0.265, error <0.001

### SM 5: Table - L-DOPA has disparate effects on forgetting rate depending on whether items were re-exposed or not

**A**

| **Variation source** | **Sum of squares** | **Mean square** | **Mean diff** (std error) | **Cohen's δ** | **F** | **p** | **$BF_{01}$** | **$H_0$ vs $H_1$** |
|---|---|---|---|---|---|---|---|---|
| **List type (*i* cf. *ii*)** | 22.293 | 22.293 | **- .798** ( .08 ) | 1.611 | 90.83 | < .001 | < . 001 | 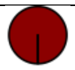 |
| *error* | *8.345* | *.245* | | | | | | |
| **Treatment** | .679 | .679 | **- .139** ( .08 ) | .280 | 2.75 | .107 | 1.4 | 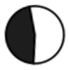 |
| *error* | *8.406* | *.247* | | | | | | |
| **Interaction** | .843 | .843 | | | 6.15 | .027 | .7 | 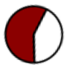 |
| *error* | *4.658* | *.137* | | | | | | |

**B**

D' ~ List Type * Dose + ( List Type + Dose | | participant)

| | **Estimate** (std error) | **t** | **df** | **p** | **$R^2$** marginal |
|---|---|---|---|---|---|
| Intercept | 1.800 ( .10 ) | 18.538 | 34.0 | <.001 | |
| List Type | .798 ( .08 ) | 9.460 | 34.0 | <.001 | |
| Dose | - .052 ( .03 ) | - 1.882 | 29.0 | .070 | .261 |
| List Type *Dose | .098 ( .04 ) | 2.302 | 32.8 | .028 | |

**L-DOPA has disparate effects on forgetting rate depending on whether items were reactivated or not.**
**A. Exploratory Parametric and Bayesian ANOVAs** show a main effect of type of encoding (List i vs List ii) and an interaction between treatment (placebo vs L-DOPA) and encoding type. $BF_{01}$ represents likelihoods of collecting our data under models that do not include the given source for variation ($H_0$ in white) compared to models that include the variation source ($H_1$ in dark). Note that $BF_{01}$ which denotes how much likelier our data are under the null are reported as opposed to $BF_{10}$ for easier interpretation. P-values are corrected for FDR using Benjamini-Hochberg procedure and accounting for 3 tests. All df = 34, 1.
**B. A planned Mixed linear model** showed that dose and encoding type together explain 26.1% of the variation in d'. As the ANOVA cannot account for dose-dependent effects, we conducted a mixed linear effects analysis with dose (mg/kg) and re-exposure (List i vs ii), and the interaction between the two, as fixed effects, and individual subjects as random effects (with slopes and intercepts). The model revealed a main effect of encoding type ($t(34) = 9.460$, $p < .001$) and a treatment * encoding type interaction ($t(32.8) = 2.302$, $p = .028$) but no evidence for a main effect of dose ($t(29) = -1.882$, $p = .070$). The model without the random effect predicted 26.1% of the variability in d'. In other words, L-DOPA enhanced performance selectively for re-activated items, while reducing performance for activated items. Note that estimates are mean-centred.

SM 6 : Table - Single dose of nocturnal L-DOPA increases time spent in slow wave sleep by 10.6%.

**Polysomnography – sleep stages**

| | **Mean** (SD) | | **Credible interval δ** | **t** | **p** (uncorrected) | **$BF_{01}$** | **$H_0$ vs $H_1$** |
|---|---|---|---|---|---|---|---|
| Time in minutes | **L-DOPA** | **Placebo** | | **df = 31** | | | |
| **Total time in bed Time** | **581.1** ( 200.3 ) | **522.9** ( 144.2 ) | [ - .269 – .650] | 365.5* | .067 | 3.6 | 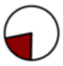 |
| **Asleep** | **363.7** ( 60.4 ) | **358.0** ( 53.8 ) | [ - .368 – .543 ] | .402 | .691 | 4.9 | 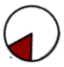 |
| **Awake After Sleep Onset** | **116.6** ( 46.6 ) | **110.5** ( 50.9 ) | [ - .272 – .636 ] | .838 | .408 | 3.8 | 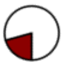 |
| **N1** | **20.0** ( 8.4 ) | **21.9** ( 8.8 ) | [ - .712 – .210 ] | - 1.160 | .255 | 2.9 | 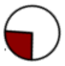 |
| **N2** | **140.9** ( 51.1 ) | **140.6** ( 58.2 ) | [ - .574 – .327 ] | - .576 | .569 | 4.5 | 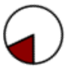 |
| **N3 (SWS)** | **132.8** ( 54.0 ) | **120.0** ( 51.3 ) | [ .116 – 1.104 ] | 2.702 | .011** | .2*** | 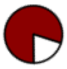 |
| **REM** | **70.0** ( 24.7 ) | **75.5** ( 25.1 ) | [ - .768 – .142 ] | - 1.426 | .164 | 2.1 | 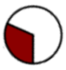 |

**Sleep stage comparisons between L-DOPA and placebo nights.**

L-DOPA increased time spent in slow wave sleep, but it did not affect light sleep (stages 1 and 2), wakefulness or total time spent asleep. Δ denotes effect size for the paired differences derived from the Bayesian posterior distribution. Where credible intervals are the 95% intervals overlapping zero denote no difference. $BF_{01}$ and $H_0$ vs $H_1$ show the probability of our data having been observed under the null (white) as opposed to the alternative (black) hypothesis. P-values are uncorrected. Note that $BF_{01}$ which denotes how much likelier our data are under the null are reported as opposed to $BF_{10}$ for easier interpretation.

*Wilcoxon test used.

** a Benjamini-Hochberg corrected p-value for slow wave sleep (N3) is 0.044, when including all sleep stages (N1, N2, N3 and REM; n tests = 4), total time in bed or time awake after sleep onset

*** corresponding $BF_{10}$ = 4.0

All errors <.055%

### SM 7 : Table – Missing data

| | Test day | | | |
|---|---|---|---|---|
| | **LDOPA** | | **Placebo** | |
| | 🏠 **0** | 📞 **1** | 🏠 **0** | 📞 **1** |
| **# of volunteers** | | | | |
| **1** | **X** | **X** | | |
| **3** | | **X** | | |
| **1** | | **X** | **X** | |
| **1** | | | | **X** |
| **2** | | | **X** | **X** |

**Summary of missing data.**

The above table provides a summary of missing data for the **SECONDARY** study. We recruited 35 volunteers to take part in the this control experiment. Two were excluded prior to dosing; due to a contraindication, and for participating in another drug trial simultaneously. Crosses (X) denote missing data points for each test session with the left column denoting the number of volunteers affected. Remaining data for each volunteer was used except where the volunteer only completed one test session (n = 3, top and bottom rows, ); two withdrew consent for personal reasons and one experienced significant nausea and vomiting as a side effect on their second testing session. Remaining data for 5 volunteers was missing partially, either due to missed phone calls or experimenter error, and one volunteer was excluded entirely as their accuracy being so low that it was below chance level – suggesting they had misunderstood the task – or due to experimenter error where wrong test versions were used.

**Main study**

Fifty-eight volunteers completed screening for the main experiment. Ten screened volunteers could not take part due to diary clashes (trial finished before they could be booked in), 2 could not take part due to incidental findings that were also contraindications revealed at screening, 3 could not take part due to existing cardiac or medicinal contraindications and a further 4 had other contraindications. 4 participants refused participation following screening (final n = 35). Further, data for one follow-up phone call (placebo, 📞 Day 5) had to be excluded due to researcher error (same set of distractors and targets were used as on 📞 Day 3). Data were partially or entirely missing for the polysomnography (PSG) for four volunteers due to technical errors at recording ($n_{PSGSWS} = 31$). One additional datapoint was excluded from spindle analyses as spindle data could not be reliably extracted ($n_{PSGSPINDLES} = 30$)

SM 8 : Table - Sleep and memory correlations on L-DOPA and placebo

| | SWS duration and D' on Day 1 | |
|---|---|---|
| **N = 31** | List i | List ii |
| **Correlations on L-DOPA** | | |
| Spearman's rho | 0.043 | 0.065 |
| *p-value* | *0.810* | *0.720* |
| **Correlations on placebo** | | |
| Spearman's rho | 0.450 | 0.320 |
| *p-value* | *0.009*<br>*P corrected = 0.012 ** | *0.071* |

**Benjamini Hochberg corrected p-value accounting for each analysis in this table*

| | SWS Spindle amplitude and D' on Day 1 | |
|---|---|---|
| **N = 31** | List i | List ii |
| **Pearson Correlations on L-DOPA** | | |
| Pearson's R | -0.026 | -0.059 |
| *p-value* | *0.891* | *0.753* |
| **Pearson Correlations on placebo** | | |
| Pearson's R | 0.234 | 0.264 |
| *p-value* | *0.151* | *0.204* |

### SM 9 : Table - L-DOPA increases spindle amplitude

**Spindle architecture during N3 - SWS**

| Measure | **Mean** (SD) **L-DOPA** | **Mean** (SD) **Placebo** | **Credible interval δ** | **t** (df = 31) | **p** (df = 31) | **p** (corrected) | **$BF_{01}$** | **$H_0$ vs $H_1$** |
|---|---|---|---|---|---|---|---|---|
| **Spindle density (# per min)** | **5.7** ( 2.2 ) | **5.8** ( 4.5 ) | [ - .351 – .313 ] | 249.0* | .992 | .992 | 5.2 | 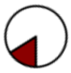 |
| **Spindle amplitude (microvolts)** | **28.9** ( 8.3 ) | **28.3** ( 8.5 ) | [ .086 – .804] | 95.0* | .002 | .008 | 0.3** | 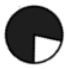 |
| **Spindle frequency (Hz)** | **13.6** ( 0.3 ) | **13.6** ( 0.3 ) | [ - .423 – .247 ] | 197.0* | .327 | .436 | 4.6 | 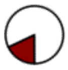 |
| **Spindle duration (sec)** | **.93** ( .05 ) | **.94** ( .06 ) | [ - .636 – .060 ] | 1.762 | .088 | .176 | 1.3 | 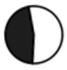 |

**Slow oscillations**

| | **Mean** (SD) **L-DOPA** | **Mean** (SD) **Placebo** | **Credible interval δ** | **t** (df = 30) | **p** (uncorrected) (df = 30) | **$BF_{01}$** | **$H_0$ vs $H_1$** |
|---|---|---|---|---|---|---|---|
| **Amplitude** | **71.8** ( 4.9 ) | **72.3** ( 4.8 ) | [ - .560 – .130] | 1.280 | .210 | 2.5 | 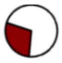 |
| **Duration (mins)** | **27.2** ( 12 ) | **29.7** ( 14 ) | [ - .162 – .517 ] | 1.083 | .287 | 3.1 | 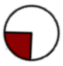 |

**NREM stage 3**
L-DOPA increased spindle amplitude. While the mean difference was small, an increase in amplitude on L-DOPA compared to the place night was seen in 25 of 31 participants, see also Fig 4a in main text.
Where credible intervals are the 95% intervals overlapping zero denote no difference. $BF_{01}$ and $H_0$ vs $H_1$ show the probability of our data having been observed under the null (white) as opposed to the alternative (black) hypothesis. P-values are uncorrected. Note that $BF_{01}$ which denotes how much likelier our data are under the null are reported as opposed to $BF_{10}$ for easier interpretation.
** corresponding $BF_{10}$ =3.6
All errors <.055%

### SM 10 : Table - Subjective sleep measures

| | Mean (SD) L-DOPA | Placebo | Credible interval δ | t | p | $BF_{01}$ | $H_0$ vs $H_1$ |
|---|---|---|---|---|---|---|---|
| | | | | df = 31 | | | |
| *St Mary's Hospital Sleep Questionnaire* | | | | | | | |
| Efficiency (%) | 75.8 ( 15.6 ) | 77.0 ( 22.9 ) | [ - .378 – .275 ] | 163* | *.156* | 5.1 | 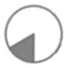 |
| Onset latency (min) | 17.1 ( 20.7 ) | 27.4 ( 31.6 ) | [ - .641 – .045 ] | 61.0* | *.060* | 1.2 | 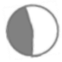 |
| Latency (min) | 390.5 ( 84.2 ) | 384.5 ( 95.1 ) | [ - .442 – .223 ] | 186* | *.952* | 4.3 | 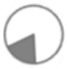 |
| Maintenance score | 3.1 ( 1.3 ) | 2.8 ( 1.2 ) | [ - .117 – .636 ] | 174* | *.270* | 2.4 | 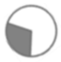 |
| Sleep satisfaction** | 62.1 ( 19.4 ) | 67.5 ( 15.4 ) | [ - .468 – .229 ] | - .694 | *.494* | 4.1 | 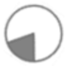 |
| Wakefulness | 31.8 ( 38.7 ) | 27.8 ( 40.7 ) | [ - .282 – .378 ] | .302 | *.765* | 5.1 | 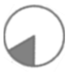 |
| *Leeds Sleep Evaluation Questionnaire* | | | | df = 34 | | | |
| Total | 4.2 ( .86 ) | 4.3 ( .96 ) | [ - .432 – .221 ] | - .668 | *.509* | 4.5 | 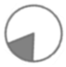 |
| GTS | 3.9 ( 1.7 ) | 3.9 ( .1.2 ) | [ - .432 – .221 ] | .109 | *.914* | 5.5 | 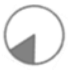 |
| QOS | 3.5 ( 1.8 ) | 3.7 ( 1.8) | [ - .432 – .221 ] | - .677 | *.503* | 4.5 | 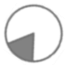 |
| AFS | 4.8 ( 1.6 ) | 4.9 ( 4.9 ) | [ - .432 – .221 ] | 304.5 | *.911* | 4.9 | 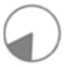 |
| BFW | 4.8 ( 1.6 ) | 4.9 ( 1.4) | [ - .432 – .221 ] | 282.5 | *.600* | 4.8 | 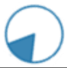 |

*Sleep: Self-evaluation*

We found no differences in self-reported measures of sleep quality. δ denotes effect size for the paired differences derived from the Bayesian posterior distribution. $BF_{01}$ and $H_0$ vs $H_1$ show the probability of our data having been observed under the null (white) as opposed to the alternative (blue) hypothesis. All p-values are uncorrected. All errors <.06.
GTS = getting to sleep, QOS = quality of sleep, AFS = awakening from sleep, BFW =behaviour following waking.
*Wilcoxon's test; ** df = 28

**Behavioural data:** Data was entirely missing for the St Mary's Hospital Sleep Questionnaire (SMHSQ) for two volunteers following the L-DOPA and one volunteer following the placebo night ($n_{SMHSQ}$ = 32). One volunteer on placebo, and two on L-DOPA, had omitted answers on the SMHSQ, so their sleep satisfaction score (SSS) could not be determined ($n_{SSS}$=29). These questionnaires were otherwise scored. The wakefulness after sleep onset score was calculated as the difference between self-reported sleep onset time and final wake up time, and self-reported sleep latency. Some volunteers reported less time between sleep onset and waking than spent asleep. For these nights, wakefulness after sleep onset was changed to 0 minutes to avoid negative values. There was no missing data for the Leeds Sleep Evaluation Questionnaire.

### SM 11 : Figure - Sleep visit timeline

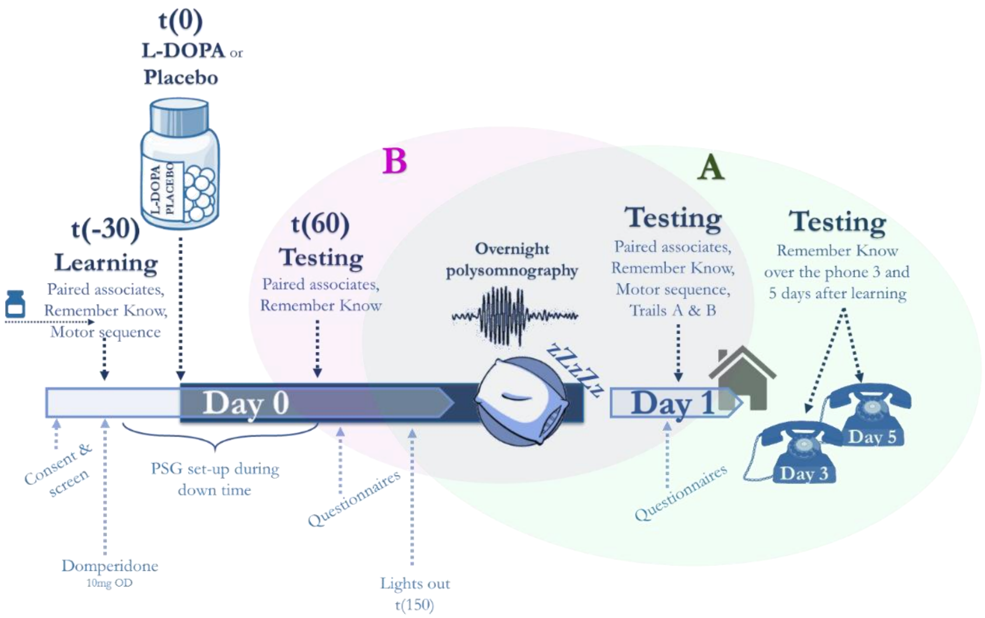

In this placebo-controlled double-blind randomised study, volunteers were initially screened over the phone for common exclusions, and then invited for three in-house visits. On the first visit the volunteers were fully screened for eligibility, demographic information was collected, and they practiced the memory tasks. Each sleep visit began with a confirmation of continued eligibility. The L-DOPA administration time is referred to as t(0). Other times are referenced to t(0) in minutes. The blue shaded area denotes time during which L-DOPA was active in the system. This study was designed to test whether L-DOPA during sleep affects either of two processes. **A**. We were interested in L-DOPAs effect on memory over long term. To assess this, we analysed sleep data and memory data for Days 1, 3, and 5 (shaded in green).

**B**. We were interested in whether L-DOPA affects memory for items that have been retrieved once before, differently from items that have not been retrieved. We tested a sub-set of items in the evening, t(60), and then re-tested these items with novel targets in the morning. We analysed the re-tested against tested items on Day 1 and contracted this with the sleep recording (shaded in purple).

Each volunteer completed two visits; on one of they received L-DOPA, on the other placebo. This figure is not to scale.

On the second visit, volunteers arrived on site in the evening and were consented and re-screened. They were provided with dinner on-site and the polysomnography was set-up. They then learnt a verbal memory task. After this they were dosed with an anti-emetic before learning other memory tests not reported here. Following the learning they were given the L-DOPA 200mg or a placebo (matched for visual appearance) in a double-blind fashion. An hour later, their memory was tested. They completed a mood questionnaire and they were given 45 minutes before their lights were switched off and polysomnography was recorded overnight. All events were calculated backward from their normal bed-time so that L-DOPA was administered 2.5h prior to sleep (and learning took place 3h prior to sleep).

A study doctor monitored participants throughout the evening with a 30s 12-lead ECG and blood pressure taken at baseline (any time prior to anti-emetic administration), immediately after domperidone was given (t(-30)), immediately prior to dosing L-DOPA (t(-05)), and 30 minutes and 60 minutes after L-DOPA (t(30), t(60)).

In the morning the volunteers were woken up at an agreed time that matched their normal sleep unless they naturally woke up earlier. They were given around 1.5h between waking up and testing during which they had a chance to shower, complete mood and sleep questionnaires, and have breakfast and refreshments. They then completed the memory tasks (~12 hours after learning) before leaving the site. 2 and 4 days later (3 and 5 days from learning), they were contacted over the phone to test their memory on the verbal memory task.

The second and third visits were identical except for treatment allocation

### SM 12 : Figure - Verbal Memory Task

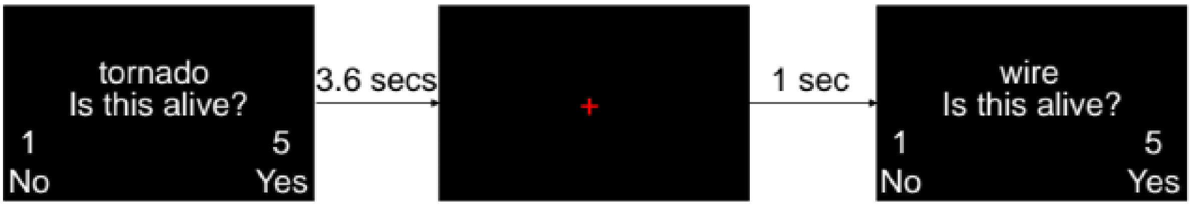

In this task, volunteers first learnt a list of 80 words presented on a computer screen in a random order, one at a time. While each word was presented, volunteers performed an incidental task where they were tasked asked to determine if the items were 'alive' or not by pressing one of two buttons on a custom-made response box (https://www.blackboxtoolkit.com). Each word was displayed once for 3.6s, whether a response was made or not, and separated by a 1s fixation cross. Words were separated into two 40-word blocks to allow a break.

In the test phase of the tasks, volunteers were shown a list of words presented individually, half of which were targets (present at learning) and half of which were distractors (not present at learning). They were asked to judge whether they had seen a word previously by judging the word as 'OLD' (target) or 'NEW' (distractor). On Days 0, 3, and 5, 20 targets and 20 distractors were shown, and on Day 1, 40 targets and distractors were shown. The same targets as were present on Day 0 were re-tested on Day 1 (i.e. they were biased as behaviourally salient by re-exposure), together with targets that had only been seen during learning and not during day 0 testing (non-salient targets). Each distractor was only seen once. The exposed and re-exposed targets were compared to study L-DOPAs effect on behaviourally "tagging" important memories with the rationale that when a word is presented a second time by a recognition test, it will be deemed to be more 'worthy' of being remembered later than a word that is presented once.

All on-screen text was displayed using Helvetica font on a Toshiba laptop. On days 0 and 1 testing was completed on a laptop, on days 3 and 5 testing was completed over the phone.

After each 'OLD' / 'NEW' judgement volunteers completed remember-know judgements on the items (*2*). The experiment was programmed in the MATLAB environment (2015b or 2017a) using the Psychophysics Toolbox V3 (*3*). The full script, standard operating procedures for administering the task, and task instructions are available from the corresponding author upon request.

### SM 13 : Figure - Secondary study timeline

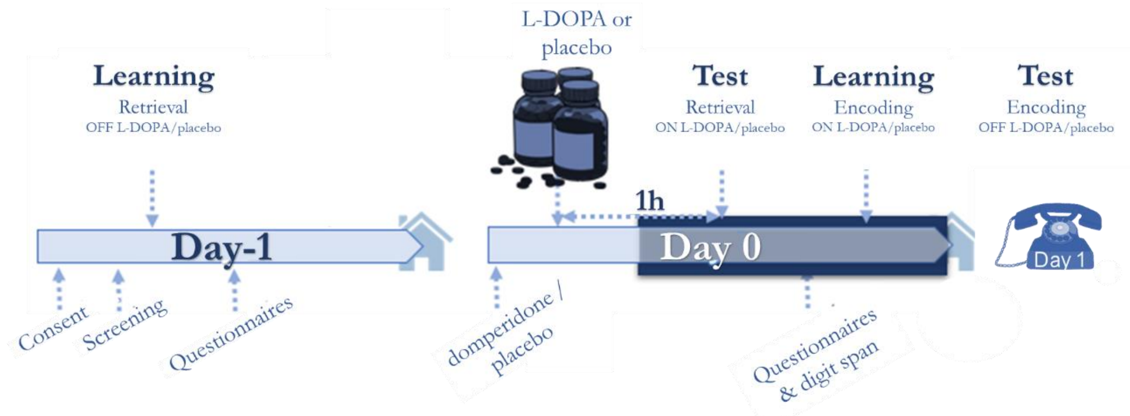

Volunteers were invited for two test sessions, each consisting of two consecutive days of testing. On Day -1 of each session the volunteers learnt an episodic verbal memory task, and their baseline performance was tested. On day 0 volunteers returned to site and they were dosed with L-DOPA or placebo before retrieval was tested. After this they learnt another episodic memory task before returning home. Their recall on this task was tested immediately and three times over the phone following a 1-day delay. Experiment 2 study aimed to test the effect of L-DOPA on retrieval and encoding.

**Day -1 :** On the day preceding dosing, or Day -1 (D-1), volunteers were consented and screened to ensure they met the eligibility criteria. They then learnt two experimental tasks; a verbal episodic memory task and a reinforcement learning task. The latter is reported elsewhere (*4, 5*). The learning phase was followed by paper assessments, and 30 minutes later a baseline memory test.

Day 0: On Day 0 (D0), volunteers returned to the test site where they were first dosed with the domperidone and their blood pressure and heart-rate was monitored. 30 minutes later, they received L-DOPA. At baseline and for 2 hours following Domperidone, volunteers' heart rate and blood pressure were monitored at 30-minute intervals as a safety procedure.

1h following drug administration retrieval was tested for both tasks. Volunteers then completed the digit span test, and they were offered to take a break before learning another episodic verbal memory task. Their learning for this test was measured immediately, and again over the phone the following day.

On one of the testing sets the volunteer received a placebo and, on another L-DOPA, otherwise the testing sets were identical. Standard operating procedures and guides for experimenters running the trial is available upon request from the corresponding authors.

### SM 14: Table - Study 2 results

**Encoding**

| | Mean (SD) | | Credible interval δ | t | p | $BF_{01}$ | $H_0$ vs $H_1$ |
|---|---|---|---|---|---|---|---|
| Day 1 | L-DOPA | Placebo | | df = 28 | | | |
| D’ | 1.257 ( .76 ) | 1.149 ( .61 ) | [ - .295 – .426 ] | - .352 | .728 | 4.6 | 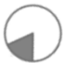 |
| Criterion | - .027 ( .35 ) | - .083 ( .39 ) | [ - .310 – .414 ] | 114* | .197 | 4.7 | 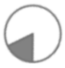 |
| Day 3 | | | | df = 25 | | | |
| D’ | .865 ( .47 ) | .710 ( .50) ) | [ - .001 – .763 ] | -2.128 | .043 | .7** | 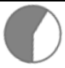 |
| Criterion | - .107 ( .40 ) | - .126 ( .46 ) | [ - .352 – .368 ] | - .040 | .968 | 4.8 | 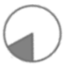 |
| Day 5 | | | | df = 26 | | | |
| D’ | .692 ( .45 ) | .592 ( .40 ) | [ - .317 – .588 ] | - .325 | .748 | 2.4 | 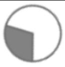 |
| Criterion | - .147 ( .38 ) | - .167 ( .33 ) | [ - .334 – .378 ] | - .148 | .884 | 4.9 | 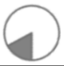 |

**2nd clinical trial Encoding: Pairwise comparisons**

* Wilcoxon test used due to non-parametric data. All samples contained zero values for paired differences. P-values for Wilcoxon tests for such data are less reliable

Only Day 1 results are reported the manuscript. However, memory was also prompted on days 3 and 5 as described in Supplementary Figure 9. The trend towards L-DOPA induced improvement in d’ at day 3 does not survive multiplicity corrections and therefore is not interpreted further in this manuscript.

**Retrieval**

| | Mean (SD) Day preceding | | Credible interval δ | t | p | $BF_{01}$ | $H_0$ vs $H_1$ |
|---|---|---|---|---|---|---|---|
| Day -1 | L-DOPA | Placebo | | df = 27 | | | |
| D’ | 2.842 ( .72) | 2.721 ( .76) | [ - .180 – .538 ] | -1.041 | .307 | 3.1 | 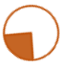 |
| Criterion | .114 ( .34 ) | .196 ( .35 ) | [ - .564 – .149 ] | -1.198 | .241 | 2.6 | 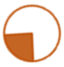 |
| Day 0 | L-DOPA | Placebo | | df = 27 | | | |
| D’ | 1.658 ( .55) | 1.609 ( .56 ) | [ - .278 – .417 ] | .393 | .698 | 4.6 | 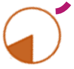 |
| Criterion | - .216 ( .39 ) | - .192 ( .47 ) | [ - .384 – .316 ] | - .224 | .968 | 4.9 | 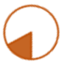 |

**Before L-DOPA / Placebo given**

**2nd study: Retrieval: Pairwise comparisons**

.

### SM 15 : Table - Demographic information

| | Encoding (experiment 2) *n= 32* | | Forgetting / re-exposure (experiment 1) *n = 35* | | | Retrieving (experiment 2) *n = 28* | |
|---|---|---|---|---|---|---|---|
| | **Mean** (SD) | **Range** | **Mean** (SD) | **Range** | | **Mean** (SD) | **Range** |
| **Age** | **71.1** (7.1) | 65 – 92 | **68.9** (3.5) | 65 – 79 | | **70.9** (6.9) | 65 – 92 |
| **Years of education** | **14.7** (3.5) | 10 – 24 | – | – | | **14.5** (3.5) | 10 – 24 |
| **MoCA** | **26.1** (3.3) | 18 – 30 | **27.5** (2.5) | 21 – 30 | | **26.1** (3.2) | 18 – 30 |
| **Height (cm)** | **170.0** (10.3) | 152 – 186 | **166.1** (7.4) | 152 – 181 | | **170.0** (10.6) | 152 – 186 |
| **Weight (kg)** | **75.2** (15.4) | 51 – 105 | **70.28** (13.0) | 48 – 94 | | **75.0** (15.6) | 51 – 105 |
| **Body mass index (kg/cm²)** | **25.8** (3.7) | 18.1 – 35.1 | **25.2** (3.2) | 18.5 – 32.7 | | **25.8** (3.7) | 18.1 – 35.1 |
| **L-DOPA concentration (mg/kg)** | **2.08** (0.44) | 1.43 – 2.95 | **2.94** (0.54) | 2.13 – 4.17 | | **2.09** (0.45) | 1.43 – 2.95 |
| **Gender (f/m)** | | **16** / **16** | | **22** / **13** | | | **14** / **14** |
| **Treatment order (L-DOPA/ Placebo first)** | | **17** / **15** | | **18** / **17** | | | **13** / **15** |
| **Blinding (accurate / inaccurate** / missing**)** | | **17** / **12** / 3 | | **21** / **13** / 1 | | | **16** / **10** / 2 |
| **Rationality Experientiality index (REI) scores** | | | | | | | |
| ***Overall rationality*** | **3.6** (0.8) | 1.5 – 5.0 | **3.5** (0.7) | 2.3 – 4.7 | | **3.7** (0.9) | 1.5 – 5.0 |
| ***Rational Engagement*** | **3.8** (0.9) | 1.0 – 5.0 | **3.4** (0.7) | 2.1 – 4.6 | | **3.8** (0.9) | 1.0 – 5.0 |
| ***Rational Ability*** | **3.5** (1.0) | 1.0 – 5.0 | **3.5** (0.7) | 2.1 – 4.7 | | **3.5** (1.0) | 1.0 – 5.0 |
| ***Overall Experientiality*** | **3.1** (0.6) | 2.0 – 4.2 | **3.4** (0.7) | 2.1 – 5.0 | | **3.1** (0.6) | 2.0 – 4.2 |
| ***Experiential Engagement*** | **2.9** (0.7) | 1.0 – 4.2 | **3.3** (0.6) | 2.2 – 5 | | **2.9** (0.8) | 1.0 – 4.2 |
| ***Experiential Ability*** | **3.3** (0.6) | 2.0 – 4.5 | **3.5** (0.7) | 1.9 – 4.9 | | **3.3** (0.6) | 2.0 – 4.3 |
| **Depression, Anxiety and Stress Scale (DASS)** | | | | | | | |
| ***Depression*** | **4.2** (4.9) | 0 – 18 | **2.1** (3.2) | 0 – 11 | | **4.0** (4.7) | 0 – 18 |
| ***Anxiety*** | **2.0** (2.4) | 0 – 11 | **1.8** (2.3) | 0 – 8 | | **2.0** (2.6) | 0 – 11 |
| ***Stress*** | **5.4** (4.1) | 0 – 14 | **6.2** (5.9) | 0 – 21 | | **5.4** (4.3) | 0 – 14 |
| **Barratt Impulsivity Scale (BIS)** | | | | | | | |
| ***Motor impulsiveness*** | **21.3** (3.7) | 14 – 29 | **21.1** (3.6) | 12 – 27 | | **20.9** (3.5) | 14 – 29 |
| ***Non-planning*** | **22.1** (5.1) | 11 – 30 | **21.7** (5.3) | 14 – 35 | | **21.6** (4.7) | 11 – 30 |
| ***Attentional*** | **14.6** (2.5) | 10 – 19 | **14.5** (3.4) | 9 – 23 | | **14.4** (2.5) | 10 – 19 |
| **Pittsburgh sleep quality index** | | – | **35** (10.4) | 20 – 59 | | – | – |
| ***Sleep efficiency*** | – | – | **1.4** (1.5) | – | – | – | – |
| ***Sleep Quality*** | – | – | **1.7** (1.6) | – | – | – | – |
| ***Daily disturbance*** | – | – | **1.4** (0.7) | – | – | – | – |

**Demographic variables.**

For the main study (middle column) BIS was missing from one volunteer. Otherwise full datasets are reported. L-DOPA concentration was calculated as drug dose (mg) / body weight (kg). In studies the encoding and retrieval study, volunteers were dosed with 150mg, in the forgetting / re-activating study with 200mg L-DOPA CR, both in the form of co-beneldopa. The volunteers for the encoding and retrieving studies are the same with a different iteration of missing data in each (see below).

### SM 16 : Table - Exclusion and inclusion criteria

**Inclusion** The inclusion criteria for Experiment 1 and 2 was the same; volunteers were over 65 years of age, native or fluent English speakers, and had normal or corrected-to-normal vision allowing them to read text on a computer screen.

**Exclusion**

Participants did not have:

- clinically significant neurological or psychiatric diagnoses as assessed by self-report and questionnaires during the screening visit.
- a diagnosis of mild cognitive impairment or dementia.
- undiagnosed skin lesions.
- sensitivity to levodopa, benserazide, or domperidone.
- lactose intolerance, galactosemia or glucose/galactose malabsorption.
- galactose intolerance.
- Lapp lactase deficiency.
- diagnosis of Huntington's Chorea.
- clinically significant intention tremor.
- known prolactin-releasing pituitary tumour (prolactinoma).
- diagnosis of glaucoma.
- a history of, or current, malignant melanoma.
- current cancer treatment.
- diagnosed unstable diabetes (people with stable type 2 diabetes diet-controlled diabetes were included)
- severe endocrine, hepatic, renal, pulmonary, or cardiac disorder.
- diagnosed electrolyte disturbances.
- known peptic ulcers.
- history of a heart-attack or prolongation of cardiac conduction intervals, or any other cardiac problems as taking domperidone increases risk of said problems.

childbearing potential or pregnancy.

Participants were also excluded if they were taking any of the following:

- dopaminergic medications.
- noradrenergic, serotonergic, or anticholinergic medications started or changed within the past 3 months.
- monoamine oxidase inhibitors (MAO-I), except if selective MAO-A or MAO-B inhibitors are given alone. MAO-A and MAO-B inhibitors given together are equivalent to non-selective MAO-inhibition and therefore volunteers taking both MAO-A and MAO-B were not included.
- cholinesterase inhibitors, except if the participant was on stable treatment (at least 3 months).
- antihypertensives containing reserpine.
- ferrous sulphate on the day of dosing.
- opioids or sympathomimetics (e.g. amphetamines, epinephrine/adrenaline) unless if the participant was able to abstain on the day of dosing.
- diazepam or other benzodiasepines, unless none taken for prior 3 days or stable dose was maintained for more than 3 months.
- ketoconazole, erythromycin or CYP3A4 inhibitors (e.g. fluoconazole, voriconazole, clarithromycin, amiodarone, telithryomycin).
- antibiotics, if taken to treat an active infection.
- hormone replacement therapy.
- anti-fungal agents (pentamidine).
- anti-malarial agents.
- antihistaminics unless stable dose for 3 months or none for 3 days prior to testing sessions.
- AIDS/HIV medications.
- Any QTc prolonging medicinal products.
- if a participant took antacids or antisecretory agents they were required not to be taken at the same time as domperidone

**Additional exclusion criteria for study 1**

For Experiment 1, we also excluded volunteers with clinically significant sleep problems in the past year. We consider a sleep disorder to be clinically significant when it results in fewer than 6 hours' sleep per night regularly, in the estimation of the volunteer, and they perceived to have impaired sleep. We also excluded sleep disorders that required intervention (including equipment or medication) likely to interfere with our protocol or people with diagnosed sleep disorders who require intervention but who are unable to or have decided not to have the intervention (e.g. people with sleep apnoea for which a mask was recommended but who could not tolerate the mask). We also excluded one volunteer for being a wheelchair user, as we could not accommodate for a carer to stay with the volunteer.

SM 17 : Figure - Secondary study results

**a) subsequent memory performance when L-DOPA given at encoding**

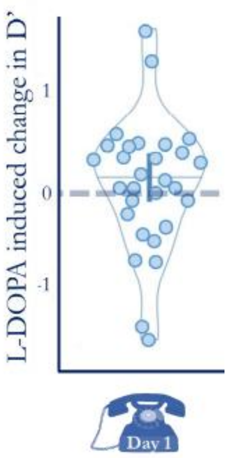

**b) memory performance when L-DOPA active during test**

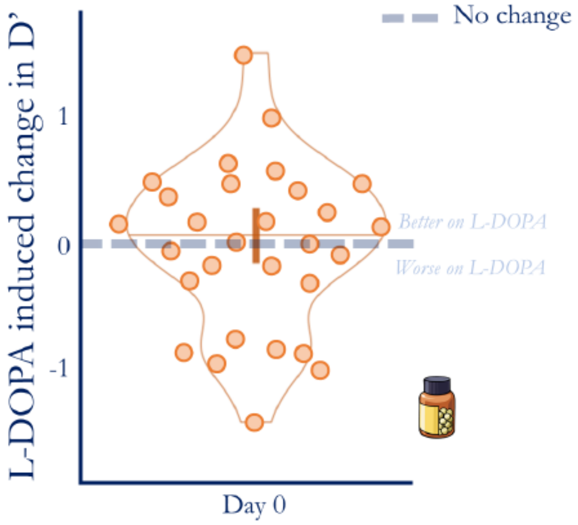

L-DOPA did not affect memory performance at a 24h delay (including a night of sleep) when given at learning (panel a) or at retrieval (panel b). Plots show paired differences between participants (L-DOPA minus placebo) for d' with minimum, median and maximum (horizontal lines) and kernel densities (vertical outlines). The dashed line denotes where there was no difference between the two visits. The data points falling below performed worse on L-DOPA compared to placebo, points above performed better.

### SM 18 : Figure - Individual Power Change

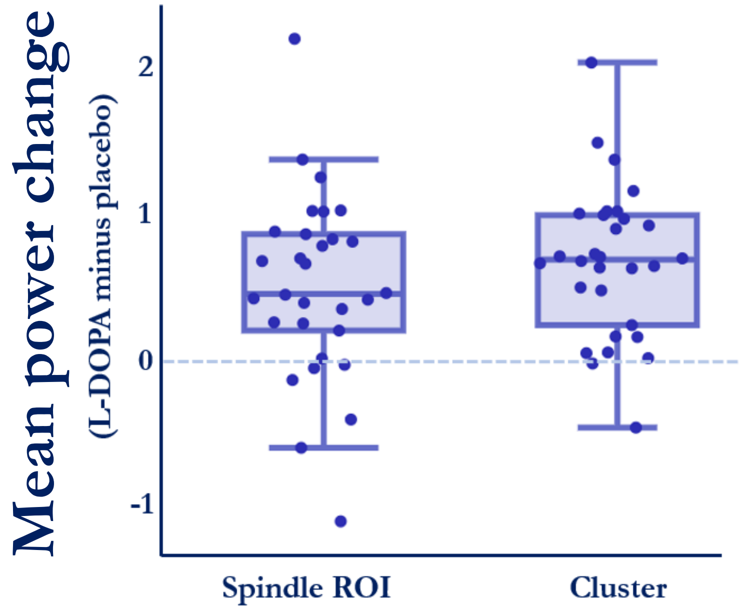

*Individual mean power change between L-DOPA and placebo conditions during slow oscillation - spindle co-occurrence events, with respect to the 2 time-frequency areas of interest: the a-priori selected spindle region of interest, 11 - 16Hz, -0.5 - 0.5s centered on the slow oscillation peak; and the primary cluster revealed by the cluster permutation method, shown in* ***Fig 4e (main text).*** *The power change for each individual was calculated by taking a mean for all power difference data points within these time-frequency spaces. While both areas were consistent across subjects, there is a very high level of consistency shown for the L-DOPA induced power increase.*

SM 18 : Diagram - CONSORT

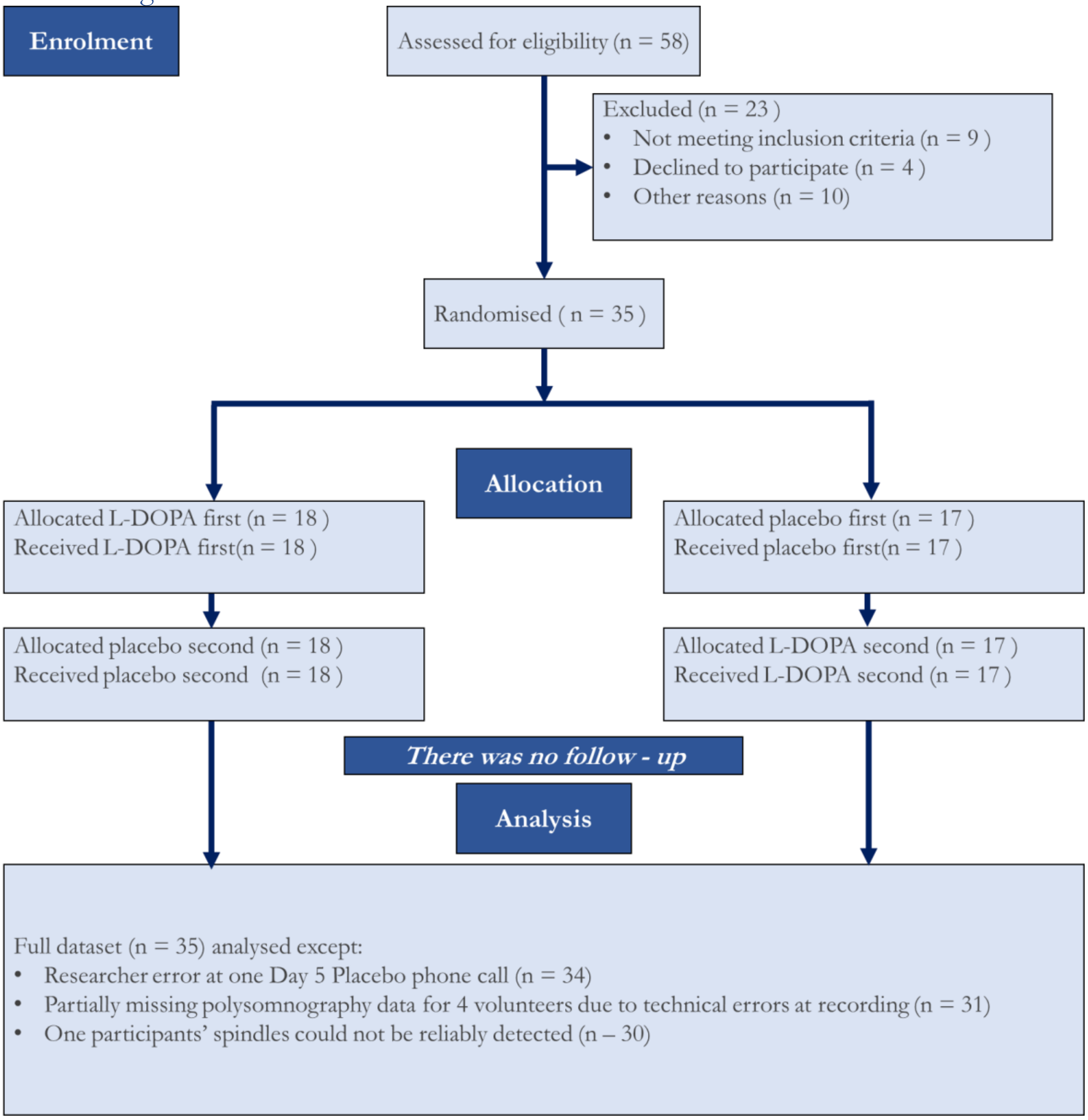

### SM - Reference list

1. K. Pearson, L. Filon, Mathematical Contributions to Theory of Evolution: IV. On the Probable Errors of Frequency Constants and on the Influence of Random Selection and Correlation. *Philosophical Transactions of the Royal Society of London. Series A, Containing Papers of a Mathematical or Physical Character* **191**, 229-311 (1898).
2. E. Tulving, Multiple memory systems and consciousness. *Hum Neurobiol* **6**, 67-80 (1987).
3. M. Kleiner, D. Brainard, D. Pelli, What's new in Psychtoolbox-3? *Perception* **36**, 14-14 (2007).
4. J. P. Grogan *et al.*, Effects of Parkinson's disease and dopamine on digit span measures of working memory. *Psychopharmacology (Berl)* **235**, 3443-3450 (2018).
5. J. P. Grogan *et al.*, Levodopa does not affect expression of reinforcement learning in older adults. *Sci Rep-Uk* **9**,  (2019).
